# Comprehensive single cell analysis of pandemic influenza A virus infection in the human airways uncovers cell-type specific host transcriptional signatures relevant for disease progression and pathogenesis

**DOI:** 10.1101/2020.04.03.014282

**Authors:** Jenna N. Kelly, Laura Laloli, Philip V’kovski, Melle Holwerda, Jasmine Portmann, Volker Thiel, Ronald Dijkman

## Abstract

Respiratory viruses, such as the 2009 pandemic strain of influenza A virus (IAV, H1N1pdm09), target cells found in the human respiratory epithelium. These cells, which form a pseudostratified epithelial layer along the airways, constitute the first line of defence against respiratory pathogens and play a crucial role in the host antiviral response. However, despite their key role in host defence, it remains unknown how distinct cell types in the respiratory epithelium respond to IAV infection and how these responses may contribute to IAV-induced pathogenesis and overall disease outcome. Here, we used single cell RNA-sequencing (scRNA-seq) to dissect the host response to IAV infection in its natural target cells. scRNA-seq was performed on human airway epithelial cell (hAEC) cultures infected with either wild-type pandemic IAV (WT) or with a mutant version of IAV (NS1_R38A_) that induced a robust innate immune response. We then characterized both the host and viral transcriptomes of more than 19,000 single cells across the 5 major cell types populating the human respiratory epithelium. For all cell types, we observed a wide spectrum of viral burden among single infected cells and a disparate host response between infected and bystander populations. Interestingly, we also identified multiple key differences in the host response to IAV among individual cell types, including high levels of pro-inflammatory cytokines and chemokines in secretory and basal cells and an important role for luminal cells in sensing and restricting incoming virus. Multiple infected cell types were shown to upregulate interferons (IFN), with type III IFNs clearly dominating the antiviral response. Transcriptional changes in genes related to cell differentiation, cell migration, and tissue repair were also identified. Strikingly, we also detected a shift in viral host cell tropism from non-ciliated cells to ciliated cells at later stages of infection and observed major changes in the cellular composition. Microscopic analysis of both WT and NS1_R38A_ virus-infected hAECs at various stages of IAV infection revealed that the transcriptional changes we observed at 18 hpi were likely driving the downstream histopathological alterations in the airway epithelium. To our knowledge, this is the first study to provide a comprehensive analysis of the cell type-specific host antiviral response to a respiratory virus infection in its natural target cells – namely, the human respiratory epithelium.

## Introduction

Respiratory viruses, including Influenza A virus (IAV), pose a significant threat to global public health and represent a major source of morbidity and mortality in humans. IAV in particular, causes not only yearly seasonal outbreaks, but also sporadic pandemics that can have devastating consequences ^1,2^. The most recent IAV pandemic, which was first detected in Mexico in 2009, was caused by the 2009 IAV H1N1 pandemic virus (H1N1pdm09) ^3^. This virus, which disproportionately affected people under the age of 65, spread quickly and ultimately led to thousands of deaths worldwide ^4^. Notably, in most people IAV causes only mild disease; however, some individuals develop more severe or even fatal disease outcomes ^5,6^. Although a number of studies have shown that IAV pathogenesis and disease severity are influenced by host innate immune and inflammatory responses, the specific host factors involved and how they shape IAV pathogenesis, remain elusive ^7,8^.

The main target cells for IAV infection and replication are airway epithelial cells ^9,10^. These cells, which form a pseudostratified layer along the human respiratory tract, are the first cells to encounter invading respiratory pathogens and play a critical role in host defence ^11^. Several distinctive cell types comprise the airway epithelium, including ciliated, secretory, goblet, and basal cells ^12^. Anchored by a collective network of adhesion molecules and cell-cell junctions, airway epithelial cells form a strong physical barrier that is impermeable to many pathogens. In addition, distinct cell types make use of unique defensive strategies to combat viral infection. For example, secretory and goblet cells secrete mucus and antimicrobial peptides onto the luminal surface of the airway epithelium, whereas ciliated cells facilitate the removal of viral and cellular debris from the respiratory tract ^12^. In addition to these extracellular defences, the host innate immune response in airway epithelial cells provides another essential layer of protection. In particular, the interferon (IFN) system coordinates the production of hundreds of different host effector proteins that (i) transform the local environment and establish an intracellular “antiviral state,” (ii) impair the propagation, spread, and transmission of viral pathogens, and (iii) shape downstream adaptive immune responses ^13–16^.

During viral infection, the IFN response is activated by the recognition of specific pathogen-associated molecular patterns (PAMPs) by one or more pattern recognition receptors (PRRs). PRRs relevant to IAV, including Toll-like receptor 3 (TLR3), Melanoma differentiation-associated protein 5 (MDA5), and retinoic acid-inducible gene I (RIG-I), are known to be expressed under both naive and IFN-stimulated conditions in the human airway epithelium ^17–20^. Upon recognition, PRRs bind to their cognate PAMPs and activate downstream signalling pathways that ultimately lead to the induction of both type I and III IFNs ^21^. Notably, type III IFNs are particularly abundant at mucosal sites and play an important role in epithelial antiviral defence, whereas type I IFNs are expressed more ubiquitously in multiple host tissues ^21^. Activation of either pathway leads to the upregulation of hundreds of IFN-stimulated genes (ISGs), as well as many pro-inflammatory cytokines and chemokines ^13–16,21^. The former interferes with viral replication directly, whereas the latter recruit and activate innate immune cells and instigate downstream adaptive immune responses.

Beyond the canonical IFN and pro-inflammatory responses, viral infection also leads to the induction of genes involved in programmed cell death (PCD), wound healing, and tissue repair. IAV infection in particular, has been shown to induce multiple forms of PCD, including both extrinsic and intrinsic apoptosis, necroptosis, and pyroptosis ^22–26^. Strikingly, recent studies have identified several PRRs that can activate both antiviral and PCD pathways following viral infection ^27,28^. Together these pathways, as well as pathways involved in host epithelium repair, play a pivotal role in dictating the severity and outcome of disease following IAV infection ^7,29–32^. However, within its natural target cells, it is currently unknown how key components of the multifaceted host response are distributed among the distinct cell types or how the latter may influence IAV disease progression and pathogenesis.

IAV has evolved various strategies to evade recognition by the host innate immune system. Many of these strategies target IFN production and as such, previous scRNA-seq studies have only been able to detect IFN in a few cells following IAV infection ^33–36^. The non-structural 1 (NS1) protein, which contains a conserved RNA-binding domain (RBD) (amino acids 1-73) and an effector domain (amino acids 74-230), is the main protein by which IAV antagonizes the host response ^37,38^. The RBD is believed to sequester viral RNA transcripts in the cell to prevent recognition of these transcripts by PRRs and avoid activation of key innate immune signalling cascades ^39^. Notably, RBD disruption at amino acid position 38 completely abrogates the dsRNA binding capacity of NS1 ^39^. Moreover, this mutation leads to attenuation of viral replication in both primary murine airway epithelial cell (mAEC) cultures and in mice _40,41_. Indeed, previous studies of mAEC cultures infected with the IAV NS1_R38A_ mutant virus (NS1_R38A_) uncovered a critical regulatory role for NS1 in both induction of and sensitivity to the host innate immune response ^40^. Therefore, despite regional differences in the cellular composition between the human and murine respiratory epithelium ^42^, and species-specific immune antagonizing effects by NS1 ^43^, this specific feature of NS1 can be utilized to exaggerate an antiviral response in order to dissect the IFN response to IAV infection in its natural target cells.

Multiple fundamental aspects of the host response to IAV infection in the human respiratory tract are unknown, including how crucial innate immune components are distributed among distinct epithelial cell types and how this distribution may influence infection outcome. Additionally, from the viral perspective, very little is known about the nature and extent of viral transcription that occurs in these cell types or how IAV infection may alter the overall composition of the respiratory epithelium. Here, we provide the first comprehensive analysis of the host response to IAV infection in its natural target cells. To mimic natural IAV infection, we infected human airway epithelial cell (hAEC) cultures with a low multiplicity of infection (MOI) of either wild-type pandemic IAV (WT) or the NS1_R38A_ mutant virus. In total, we characterized the individual host and viral transcriptomes for more than 19,000 single cells across the 5 major hAEC cell types in mock, WT, and NS1_R38A_ infected cultures. We observed a large heterogeneity in viral burden and disparate host response among virus-infected and bystander cell populations accompanied with a dynamic change in the cellular composition in both the WT and NS1_R38A_ infected cultures. This revealed that infected cells are the main producers of IFNs, with a dominant role for IFN lambda. Furthermore, we observed transcriptional changes among genes associated with inflammasome activation, cell death, wound healing, and tissue repair, that are likely responsible for the observed downstream phenotypic changes in airway epithelial cell architecture during later stages of infection. Collectively, these results provide a comprehensive overview of the complex antiviral response to IAV infection and the associated viral pathogenesis at the natural site of infection, namely the human respiratory epithelium.

## Material and Methods

### Cell lines

The human embryonal kidney cell line 293LTV (Cellbiolabs; LTV-100) was maintained in Dulbecco’s Modified Eagle Medium supplemented with GlutaMAX (Gibco), 1mM sodium pyruvate (Gibco), 10% heat-inactivated fetal bovine serum (FBS; Thermo Fisher Scientific), 100 µg/ml Streptomycin (Gibco), 100 IU/ml Penicillin (Gibco) and 0.1 mM MEM Non-Essential Amino Acids (Gibco). The Madin-Darby Canine Kidney II (MDCK-II) cell line was maintained in Eagle’s Minimum Essential Medium (EMEM, Gibco), supplemented with 5% heat-inactivated FBS (Thermo Fisher Scientific), 100 µg/ml Streptomycin and 100 IU/ml Penicillin (Gibco). All cell lines were maintained at 37°C in a humidified incubator with 5% CO_2_.

### Primary human airway epithelial cell (hAEC) cultures

Primary human bronchial cells were isolated from patients (>18 years old) undergoing bronchoscopy or pulmonary resection at the Cantonal Hospital in St. Gallen, Switzerland, in accordance with our ethical approval (EKSG 11/044, EKSG 11/103 and KEK-BE 302/2015). Isolation and establishment of well-differentiated primary human airway epithelial cell cultures was performed as previously described ^44^. The hAEC cultures were allowed to differentiate for at least four weeks prior to use.

### Recombinant Influenza A virus

The Influenza A/Hamburg/4/2009 (H1N1pdm09) virus strain in the pHW2000 reverse genetic backbone was kindly provided by Martin Schwemmle, University of Freiburg, Germany, and was used as template to generate the Influenza A H1N1pdm09NS1_R38A_ virus mutant using site-directed mutagenesis _45_. Both H1N1pdm09 (WT) and H1N1pdm09NS1_R38A_ (NS1_R38A_) viruses were rescued by transfecting 1 µg of each of the eight individual genomic segments into co-cultures of 293LTV and MDCK-II cells using lipofectamine 2000 according manufacturer instructions (Thermo Fisher Scientific). After 6 hours the maintenance medium was exchanged to infection medium (iMEM), which is composed of Eagle’s Minimum Essential Medium (EMEM), supplemented with 0,5% of BSA (Sigma-Aldrich), 100 µg/ml Streptomycin and 100 IU/ml Penicillin (Gibco) and 1 µg/mL Bovine pancreas-isolated acetylated trypsin (Sigma-Aldrich) and 15 mM HEPES. Forty-eight hours post-transfection virus containing supernatant was cleared from cell debris through centrifugation for 5 minutes at 500x *rcf* before aliquoting and storage at -80°C. Working stocks were prepared by propagating the rescued virus onto MDCK-II cells for 72 hours in iMEM after which the supernatant was clarified from cellular debris before aliquots were stored at -80°C. The viral titer was either determined by TCID50 or by Focus Forming Unit (FFU) unit assay on MDCK-II cells as described previously ^46,47^.

### Single cell RNA-sequencing of hAEC cultures

hAEC cultures from 2 different human donors were inoculated in duplicate with 10,000 TCID_50_ of either the WT or NS1_R38A_ virus or Hank’s Balanced Salt Solution (HBSS) as mock (untreated) control and incubated for 1 hour at 37°C in a humidified incubator with 5% CO2. Afterwards inoculum was removed, and the apical surface was washed three times with HBSS, after which the cells were incubated for an additional 17 hours at 37°C in a humidified incubator with 5% CO2. For each condition one of the duplicate samples was fixed in 4% formalin solution for later immunofluorescence analysis. The apical surface of the remaining inserts was washed four times with 200 µL of HBSS followed by a final washing step of both the apical and basolateral surface with 200 and 500 µL of HBSS, respectively. Cells were dissociated from the Transwell® insert by adding 200 and 500 µL of TrypLE (Thermo Fisher Scientific) to apical and basolateral compartment and an incubation step of 10 minutes at 37°C in a humidified incubator with 5% CO_2_. This was followed by a gentle disruption of the cell layer through pipetting using a large bore-size pipette tip, and an additional incubation of 20 minutes at 37°C in a humidified incubator with 5% CO_2_. Dissociated cells were transferred into 800 µL wash solution, which is composed of Air-Liquid Interface (ALI) medium supplemented with 0.1% Pluronic (Thermo Fisher Scientific), and the remaining clumps were gently disrupted through pipetting using a large bore-size pipette tip. Next three cycles of centrifugation for 5 minutes at 250x *rcf* and resuspension in 1 mL washing solution were performed. Afterward the cells were resuspended in 300 µL washing solution and the cell number, cell viability and cell size were assessed with trypan blue on a Countess II (Thermo Fisher Scientific). The single cell partitioning was performed on a Chromium Controller (10x Genomics) using the Chromium Single Cell 3’ Reagent Kit (version 2, 10x Genomics) according to manufacture protocol. The obtained partitions were further processed using the Chromium Single Cell 3’ Reagent Kit (version 2, 10x Genomics) to generate Nextera XT sequencing libraries that were sequenced on a HiSeq3000 (Illumina), using a single flow cell lane for each library.

### Computational analysis scRNA-seq data

The raw sequencing data was processed using the CellRanger software package (10x Genomics, version 2.1.1) with a concatenation of the human genome CRCh38 and the viral H1N1pdm09 strain as reference sequence. The resulting unique molecule identifier (UMI) count matrix of each individual sample was pre-processed, filtered individually in Seurat (v2.3.4) by plotting global distribution of gene, UMI, and mitochondrial counts per cell for each sample ^48^. Cell partitions that expressed genes in fewer than 5 cells, along with those that expressed fewer than 1000 genes or for which the total mitochondrial gene expression was greater than 30% were removed. Following the preliminary analyses and filtering the data of the 3 different conditions (mock, WT, and NS1_R38A_) was merged with its biological counterpart (donor 1904 or 2405) prior to data scaling, normalization and regressing out unwanted sources of variation (number of UMI’s, mitochondrial content, cell cycle phase and proportion of viral mRNAs) prior to integrating all 3 different conditions via canonical correlation analysis (CCA). The proportion of viral mRNA found in a cell was inferred from the amount of unique molecular identifiers (UMI) that aligned with viral segments in that cell. This number was then divided by the total UMI count (cellular and viral mRNAs) in the same cell to give proportion per cell. Because of potential ambient viral RNA contamination in neighbouring cell partitions in the WT and NS1_R38A_ virus-infected conditions we categorized cells as either “virus-infected” or “bystander” when the proportion of viral mRNA was above or below a threshold of 0.05, respectively. For the computation of the viral heterogeneity, fraction of missing genes and relative gene expression of influenza virus in hAEC cultures we modified previous published scripts ^35^. For cell type annotation, the resulting integrated dataset was used for unsupervised graph-based clustering was used to annotate the different cell types in mock, and WT and NS1_R38A_ virus-infected hAEC cultures using both cluster-specific marker genes and well-known canonical marker genes to match identified clusters with specific cell types found in the respiratory epithelium. Further downstream analysis, such as differential gene expression, pathway enrichment analysis and data visualization was performed with a variety of R-packages ^48–51^. Calculations were performed on UBELIX (http://www.id.unibe.ch/hpc), the High Performance Computing (HPC) cluster at the University of Bern.

### Complete Influenza virus genome sequencing

Viral RNA was extracted from 10,000 TCID_50_ WT and NS1_R38A_ virus containing inoculum using the QIAamp Viral RNA mini kit (Qiagen), according to the manufactures protocol. The viral genomic segments were amplified using the SuperScript IV One-Step RT-PCR system (Thermo Fisher) according to the previously described M-RTPCR protocol ^52^. Amplified PCR products were analysed on a 2100 bioanalyzer system (Agilent) using a High Sensitivity DNA chip according to the manufactures guidelines. The sequencing libraries of the individual samples were prepared using the Oxford Nanopore Technology (ONT) ligation sequencing kit (SQK-LSK109) in combination with the native barcoding kit (EXP-NBD104). The barcoded samples were pooled together and loaded on a MinION flowcell (ONT, R9.4) mounted on a MinION MK1b device and sequenced using MinKNOW software (v2.1), according to manufacture protocols. The raw squiggle data was processed and demultiplexed using the Albacore basecaller (v2.3.4). Reads from the inoculum virus samples were then aligned against the reverse genetic plasmid-based Influenza A/Hamburg/4/2009 reference sequence using minimap2 (v2.11) after which nucleotide variants were called with Nanopolish and translated into a new consensus sequence (v0.11.1) ^53^. Sequencing depth for the genomic segments in each sample was analysed with Samtools (v1.8) ^54^. Calculations were performed on UBELIX (http://www.id.unibe.ch/hpc), the High-Performance Computing (HPC) cluster at the University of Bern.

### Immunofluorescence

The hAEC cultures were fixed and stained for immunofluorescence as previously described ^44^. The mouse monoclonal antibody directed against the Influenza A Virus NP Protein (clone C43; ab128193, Abcam) and polyclonal rabbit anti-ZO1 (Tight junctions; 61-7300, Thermo Fisher Scientific) were used as primary antibodies. Alexa Fluor® 488-labeled donkey anti-mouse IgG (H+L), Alexa Fluor® 647-labeled donkey anti-Rabbit IgG (H+L) (Jackson Immunoresearch) were applied as secondary antibodies. The Cy3-conjugated mouse monoclonal anti-beta tubulin antibody (TUB2.1; ab11309, Abcam) was applied as a tertiary antibody to visualize the cilia. All samples were counterstained with DAPI (4’,6-diamidino-2-phenylindole; Invitrogen) to visualize nuclei. The immunostained inserts were mounted on Colorforst Plus microscopy slides (Thermo Fisher Scientific) in ProLong Diamond antifade mounting medium (Thermo Fisher Scientific) and overlaid with 0.17 mm high precision coverslips (Marienfeld). Imaging was performed by acquiring 200 nm stacks over the entire thickness of the sample using a DeltaVision Elite High-Resolution imaging system (GE Healthcare Life Sciences) using a step size of 0.2 µm with a 60x/1.42 oil objective. Images were deconvolved and cropped using the integrated softWoRx software package and processed using Fiji software package ^55^. Brightness and contrast were adjusted identically for each condition and their corresponding control. For quantification, the TJP1/ZO1 marker was used to segment cells using the Interactive Marker-controlled Watershed plugin ^56^. The subsequent mask was then used to measure cell sizes as well as tight junction intensity in a 15-pixel band corresponding to the cell periphery and based on the initial mask. Cells at the edge of the field of view were excluded from the analysis.

### Data

Single cell transcriptome data will be deposited in an open-access public repository, and scripts used for analysis and figure generation will be become available at Github upon publication.

## Results

### Single-cell RNA sequencing of pandemic IAV-infected human airway epithelial cells

To define the host response to pandemic IAV infection in its natural target cells, we infected primary human airway epithelial cell (hAEC) cultures with pandemic IAV at a multiplicity of infection (MOI) of 0.03 and then profiled the transcriptomes of uninfected cells as well as cells harvested 18 hours post-infection (hpi) using single-cell RNA sequencing (scRNA-seq) **(Fig. 1a**) ^57^. hAEC cultures derived from two distinct biological donors were infected with either wild-type pandemic IAV (WT) or a NS1 mutant virus (NS1_R38A_) with abrogated dsRNA binding capacity _39-27_. Prior to infection, both WT and NS1_R38A_ were rescued from cloned DNA and minimally passaged on MDCK-II cells and then whole genome amplicon sequencing was used to confirm that no genetic changes were introduced following viral passaging (**Supp. Table 1**). Quantification of the apical viral yield at 18 hpi revealed that the NS1_R38A_ infectious viral progeny were two-fold lower than the WT infectious viral progeny; however, their viral RNA yields were comparable **(Fig. 1b**). The former observation, which has been reported previously, suggests that the mutation in NS1_R38A_ negatively influences the production of infectious viral progeny _37,38_. Notably, we also performed whole genome amplicon sequencing on the WT and NS1_R38A_ virus-infected hAECs at 18 hpi and found that no additional mutations were introduced during multi-cycle replication (**Supp. Table 1**). As such, any discrepancies in the host response between WT and NS1_R38A_ virus-infected hAECs are due to the single non-synonymous R38A mutation in the NS1 gene.

**Figure 01.**
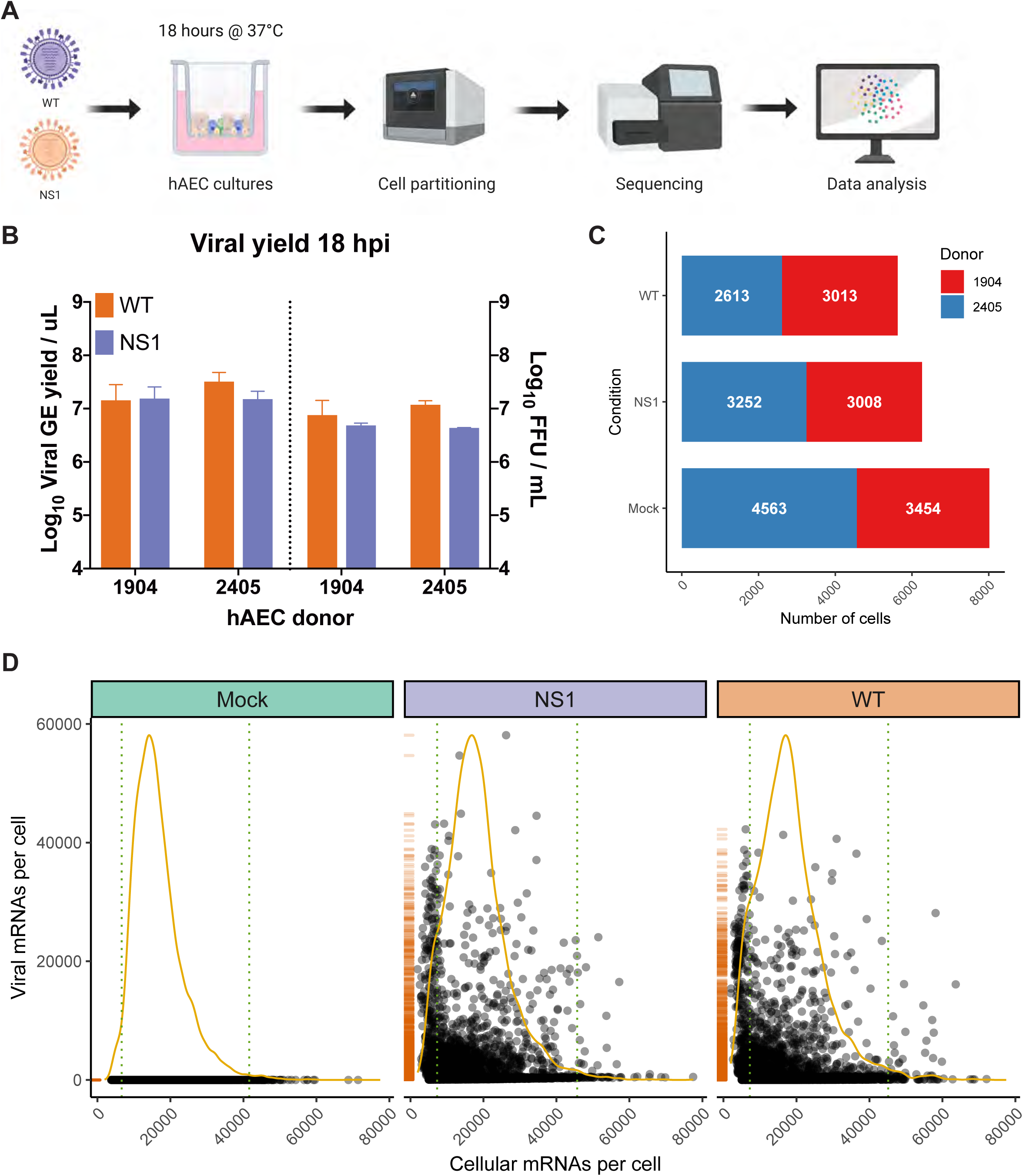
Overview of scRNA-seq workflow and evaluation of captured single cells. **A)** Overview of the experimental workflow for scRNA-seq analysis. Primary well-differentiated hAEC cultures were infected with either wild-type pandemic IAV (WT) or a mutant version of IAV (NS1_R38A_) that no longer antagonizes the host antiviral response. Cells were harvested 18 hours post-infection (hpi) and then partitioned into single cells using the Chromium Controller (10x Genomics). mRNA from single cells was subsequently reverse transcribed into cDNA and cDNA libraries were prepared and sequenced on the Illumina platform. The Cell Ranger pipeline (10x Genomics) was used to align and count both host and viral reads in individual cells and a variety of software packages were employed for downstream data analysis. **B)** Quantification of apical viral yield at 18 hpi for hAEC cultures that were derived from 2 distinct biological donors (1904 and 2405). Apical viral yield was determined for both WT (orange) and NS1_R38A_ (purple) virus-infected conditions and is given as the viral RNA yield (genome equivalents/µL; left y-axis) and viral titer (Focus Forming Units (FFU)/mL; right y-axis). **C)** Stacked bar graph illustrating the number of single cells captured for analysis. Shown for mock, WT, and NS1_R38A_ virus-infected hAEC cultures for each biological donor (donor 1904 shown in red, donor 2405 shown in blue). **D)** Scatter plot displaying the total number of host and viral mRNAs per cell for each condition (mock in green, NS1_R38A_ in purple, WT in orange). Each point in the graph represents an individual cell. The global distribution for host mRNA is shown as a yellow line, whereas the global distribution of viral mRNA is depicted as an orange rug plot.

Following IAV infection, we collected approximately 300,000 cells from our mock, WT, and NS1_R38A_ virus-infected hAEC cultures for each biological donor. Cells were then partitioned for cDNA synthesis and barcoded using the Chromium controller system (10x Genomics), followed by library preparation, sequencing (Illumina), and computational identification of individual cells **(Fig. 1a**). We captured a total of 20,282 single cells, 19,903 of which remained following the removal of cells that expressed an unusually low or high number of genes or an abnormally high amount of mitochondrial RNA (**Supp. Fig. 1a**). These 19,903 cells were comprised of 8,017 mock cells, 5,626 WT cells, and 6,260 NS1_R38A_ cells **(Fig. 1c**). Of note, since our partitioning input was 10,000 cells per condition, our recovery rate was consistent with the previously reported rate of 50-65% ^58^.

Global analysis of both host and viral transcriptomes in all 19,903 cells revealed that in each condition the host mRNA transcripts displayed an expected binomial distribution **(Fig. 1d**). Moreover, the viral mRNA transcripts were only detected in WT or NS1_R38A_ virus-infected conditions **(Fig. 1d**). We found that the proportion of viral mRNA per cell varied considerably among cells in primary hAEC cultures, which is similar to previous studies in IAV-infected A549 cells or mice ^35,59,60^.

### Identification of infected and bystander cells in IAV-infected hAEC cultures

When a population of cells is infected with a virus, not all cells in this population become truly infected. Instead, some cells, referred to as bystanders, are exposed to the virus but remain uninfected. Since infected and bystander cells have previously been shown to respond to viral infection in distinct ways, it is important to demarcate and compare these two populations. Thus, similar to previous studies, we applied a threshold that categorized cells as infected or bystander and removed any empty partitions containing displaced viral mRNAs ^35,36^. The latter occasionally occurs due to the nature of droplet-based single cell sequencing, whereby highly abundant transcripts, such as lysis-derived host mRNAs and viral mRNAs, may “leak” into neighbouring single cell partitions ^35,36,58^. Using this approach, we classified 1625 and 1701 cells as “infected” in the WT and NS1_R38A_ virus-infected samples, respectively **(Fig. 2A**). For each sample, this represents approximately 30% of all cells exposed to the virus and suggests that WT and NS1_R38A_ had infected a similar proportion of cells at 18 hpi. The latter was confirmed using immunofluorescence in a parallel experiment **(Fig. 2B**). The remaining cells in the WT and NS1_R38A_ virus-infected hAEC cultures were categorized as bystanders (4001 and 4559, respectively), whereas all cells in the mock hAEC cultures were categorized as unexposed.

**Figure 02.**
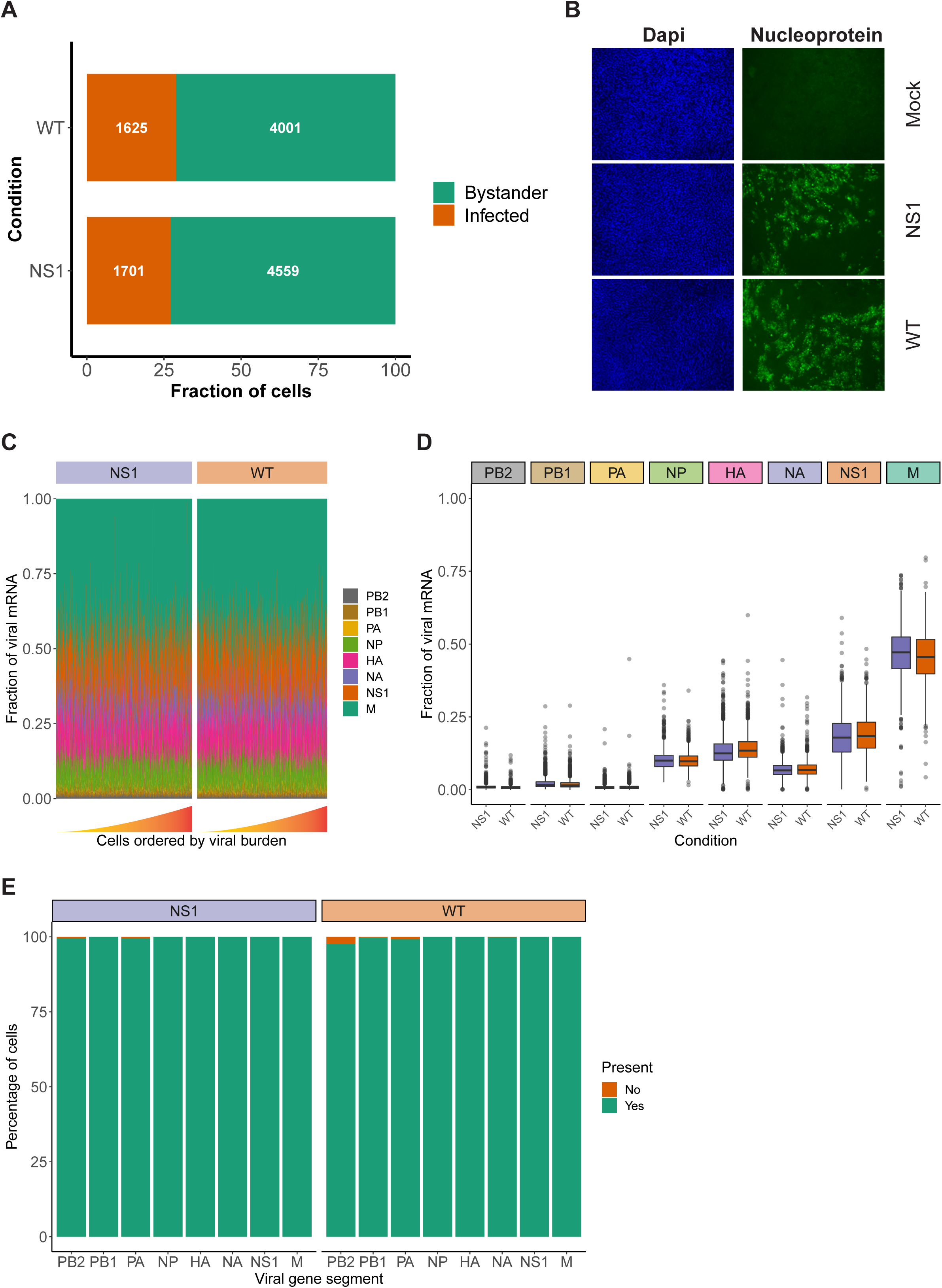
Assessment of viral characteristics in WT and NS1_R38A_ virus-infected cells. **A)** Bar graph illustrating the fraction of infected (orange) and bystander (green) cells present in both WT and NS1_R38A_ virus-infected hAEC cultures. **B)** Immunofluorescence images showing mock, WT, and NS1_R38A_ virus-infected hAEC cultures stained with DAPI (nuclei in blue; panel on left) and for the viral nucleoprotein antigen (nucleoprotein shown in green; panel on right). Representative images for each condition are shown at 18 hpi. **C)** Relative fraction of expression of viral mRNA per cell in both WT and NS1_R38A_ conditions for each IAV gene segment (only infected cells are included). Cells are ordered by increasing viral burden (from left to right) along the x-axis. **D)** Box plot summarizing the relative fraction of viral mRNA for each IAV gene segment for both WT (orange) and NS1_R38A_ (purple) conditions. **E)** Bar plot showing the percentage of infected cells expressing each of the 8 viral gene segments for both WT and NS1_R38A_ conditions. The percentage of cells whereby a specific IAV gene segment is present (green) or absent (orange) is shown for each viral gene segment (x-axis).

Since NS1 plays a major role in IAV replication we also assessed whether the R38A mutation may influence the relative expression of the different viral mRNA segments. Our analysis showed that R38A did not alter the relative expression of the viral mRNA segments and that both WT and NS1_R38A_ virus-infected hAEC cultures displayed a similar viral mRNA segment ratio with the order M > NS >> HA > NP > NA >> PB2 ∼ PB1 ∼ PA **(Fig. 2C,D**). Interestingly, this order is distinct from the order that was previously observed in IAV-infected A549 cells (M > NS >> NP > NA > HA >> PB2 ∼ PB1 ∼ PA) ^35^. Moreover, in both WT and NS1_R38A_ virus-infected hAEC cultures we found that the vast majority of infected cells express all the 8 viral mRNA segments in infected cells **(Fig. 2E**). The latter is in stark contrast to previous studies on IAV using prototypic viral strains and indicates that infection in a more natural *in vivo*-like environment with a contemporary strain of IAV exhibits distinct viral properties ^35^.

### Dynamic changes in cell composition occur in the respiratory epithelium following IAV infection

As mentioned previously, the respiratory epithelium is comprised of several specialized cell types that likely respond to IAV infection in distinct ways. To annotate these cell types and identify potential cell type-specific host and/or viral responses in mock, WT, and NS1_R38A_ virus-infected hAEC cultures, we performed unsupervised graph-based clustering on the integrated dataset using Seurat **(Fig. 3A**). We then used both cluster-specific marker genes and well-described canonical marker genes to match identified clusters with specific cell types found in the respiratory epithelium **(Fig. 3B**). In all hAEC cultures, we identified 5 distinct clusters, 4 of which corresponded to the well-known basal, secretory, goblet, and ciliated cell populations, and 1 cluster that corresponded to the recently described preciliated cell type **(Fig. 3C**). We also detected several cells expressing high levels of FOXI1, a recently defined marker for a rare group of cells called ionocytes **(Fig. 3B**). However, because these cells are rare, and since we only detected a few, we chose to exclude them from our subsequent analyses ^59,61^. Finally, we observed 1 small satellite cluster that was comprised mainly of cells from our WT and NS1_R38A_ virus-infected hAEC cultures. Careful inspection of this cluster revealed that the majority of cells found in it expressed very high levels of viral transcripts, suggesting that the host cell transcript levels may have been too low to classify these cells as a particular type. Cells in this cluster were thus categorized as “undefined” in our dataset **(Fig. 3C**). Notably, the observed cell types are consistent with previous scRNA-seq studies and indicate that our hAEC model recapitulates the respiratory epithelium *in vivo* ^42,59,61^.

**Figure 03.**
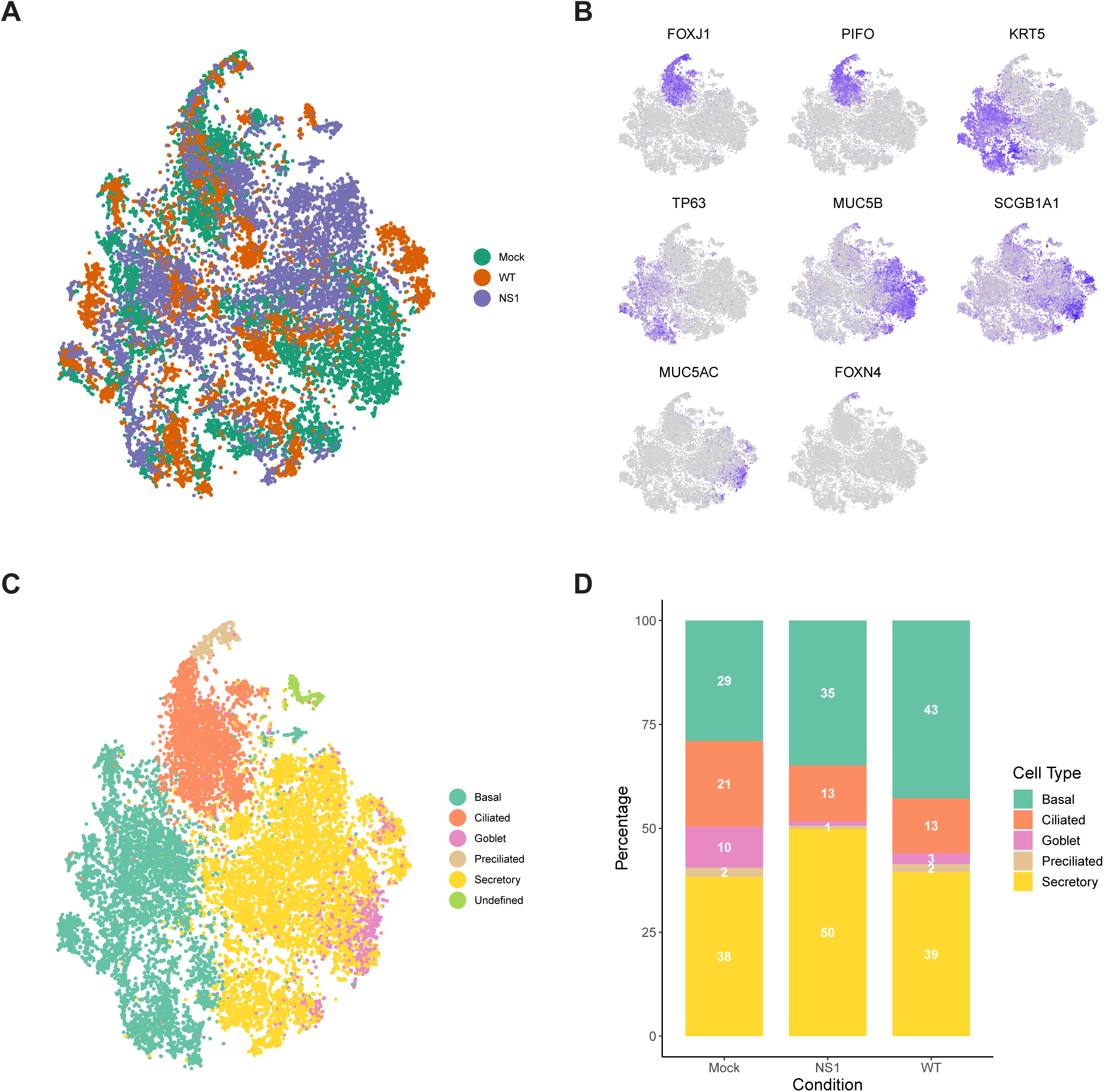
Dynamic changes in cellular composition following pandemic IAV infection. **A)** t-distributed stochastic neighbour embedding (t-SNE) visualization of the scRNA-seq data for all single cells in the mock (green), WT (orange), and NS1_R38A_ (purple) conditions following integration of the datasets in Seurat. **B)** t-SNE plots illustrating the expression patterns of several canonical airway epithelial cell type-specific markers (purple). The t-SNE plots shown in Figures A-C are presented in the same spatial orientation (i.e. the location of cells expressing the canonical markers in Figure B corresponds to the location of the specific cell types in Figure C). **C)** t-SNE visualization of the major cell types in primary well-differentiated hAEC cultures. Individual cell types were annotated using a combination of unsupervised graph-based clustering in Seurat and expression analysis of canonical cell type-specific markers. **D)** Stacked bar graph showing the relative percentage of each cell type in mock, WT, and NS1_R38A_ virus-infected hAEC cultures at 18 hpi.

To determine whether any changes in the cellular composition occurred during viral infection, we compared the relative proportion of distinct cell types found in mock hAEC samples to those found in WT and NS1_R38A_ virus-infected hAEC samples. Interestingly, compared to uninfected hAEC cultures, we observed a pronounced reduction in the ciliated and goblet cell populations for both WT and NS1_R38A_ virus-infected hAEC cultures **(Fig. 3D**). We also found an increase in the basal cell population in both virus-infected samples; however, this increase was more pronounced (34% versus 42%) in the WT sample than the NS1_R38A_ sample **(Fig. 3D**). Lastly, compared to uninfected hAEC sample, we detected an increase in secretory cells in the NS1_R38A_ virus-infected hAEC sample only **(Fig. 3D**). Since this increase is not observed in the WT sample, it is possible that WT’s ability to counteract the host antiviral response is responsible. Taken together, our results indicate that the cellular composition of the human respiratory epithelium undergoes dynamic changes during IAV infection.

### Ciliated cells become infected at later time points during pandemic IAV infection

Human-associated influenza viruses’ have a predominant affinity for non-ciliated cells, such as secretory cells, and therefore it is intriguing that we observed a decline in ciliated cells instead ^62,63^. To determine if this decline or other changes in the cellular composition correlated with cell tropism and/or viral burden, we first established the viral distribution and burden per cell type in WT and NS1_R38A_ virus-infected cultures. We categorized individual cells by both infection status and cell type to identify the proportion of infected cells for each type **(Fig. 4A**). For both WT and NS1_R38A_ virus-infected cultures, we identified infected cells in all distinct cell types; however, the majority of infected cells were found in the secretory and ciliated cell populations **(Fig. 4B**). A small proportion of basal cells were infected in both WT and NS1_R38A_ virus-infected samples, whereas a relatively large proportion of goblet and preciliated cells were infected in both conditions **(Fig. 4A**).

**Figure 04.**
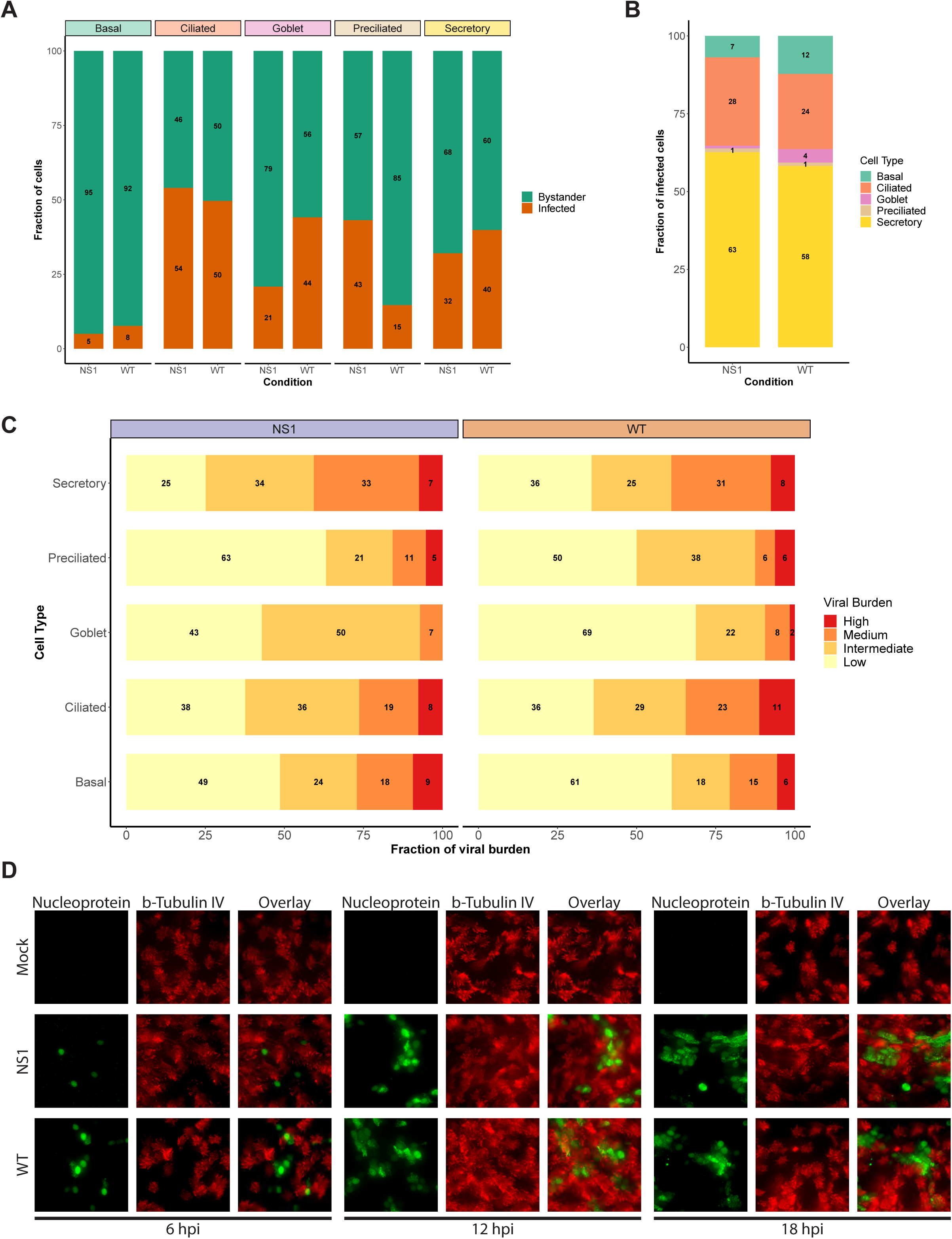
Shift in viral host cell tropism at later stages of pandemic IAV infection. **A)** Stacked bar graph displaying the percentage of infected (orange) and bystander (green) cells per cell type for both WT and NS1_R38A_ virus-infected hAEC cultures at 18 hpi. **B)** Bar graph showing the percentage of infected cells broken down by cell type for both WT and NS1_R38A_ conditions. **C)** Graph showing the relative viral burden (x-axis; low, intermediate, medium, or high) among infected cells in each cell type (y-axis) for both WT and NS1_R38A_ conditions. **D)** Immunofluorescence staining showing viral host cell tropism at 6, 12, and 18 hpi for mock, WT, and NS1_R38A_ hAEC cultures. For each time point, the viral antigen is shown in green (nucleoprotein, left panel), the ciliated cells are shown in red (ß-tubulin IV; middle panel) and the overlay is shown in the right panel.

To further investigate viral distribution, we then sub-categorized infected cells from each cell type by viral burden. To this end, infected cells were grouped to those with a low viral burden (2-10% viral mRNA), an intermediate viral burden (10-25% viral mRNA), a medium viral burden (25-50% viral mRNA), or a high viral burden (≥50%). Interestingly, we detected the highest number of infected cells with either a high or medium viral burden in the secretory cell populations of both the WT and NS1_R38A_ virus-infected hAEC cultures **(Fig. 4C**). In contrast, the goblet and preciliated populations contained the lowest number of infected cells with either a high or medium viral burden; however, this could be due to the small size of these populations. Compared to infected cells in the secretory cell population of both WT and NS1_R38A_ virus-infected samples, the overall viral burden was lowest in the basal cell population, whereas it was intermediate in the ciliated cell population **(Fig. 4C**). These results likely indicate that distinct cell types become infected at various times throughout IAV infection and/or that certain cell types may be more permissive to IAV infection than others.

It is important to note that since the average replication cycle of IAV is approximately 6 - 8 hours ^64,65^, and because we performed single cell RNA sequencing at 18 hpi, we could not discriminate between cells that became infected early on from cells that became infected at later time points during infection. Thus, to elucidate whether distinct cell types became infected at different time points throughout infection, we monitored in a parallel experiment WT and NS1_R38A_ cell tropism in virus-infected hAEC cultures at 6, 12, and 18 hpi via immunofluorescence analysis. In line with previous reports, we observed that non-ciliated cells were the predominant initial target cell population for both WT and NS1_R38A_ viruses; however, beyond 12 hpi, we detected positive IAV-antigen signal that occasionally overlapped with ciliated cell markers (e.g. beta-tubulin IV) **(Fig. 4D**). The latter indicates that distinct cell types become infected over the course of IAV infection and supports our aforementioned finding that secretory (non-ciliated) cells harbour the highest viral burden in WT and NS1_R38A_ virus-infected hAEC cultures. This indicates our scRNA-seq dataset includes both major and minor target cell type populations that become infected during the first 18 hours of a pandemic IAV infection.

### Disparate global host response among infected and bystander cell populations

We next sought to elucidate the global host response to pandemic IAV infection in its natural target cells. Following the categorization of individual cells into unique populations based on their infection status (unexposed, infected, bystander), cell type (ciliated, secretory, basal, goblet, preciliated), and viral burden, cells were placed in one of four main subsets for analysis: WT infected, WT bystander, NS1_R38A_ infected, or NS1_R38A_ bystander (**Supp. Fig. 02**). We then performed both global and cell type-specific differential gene expression analysis between each subset and the equivalent cells in the mock (unexposed) hAEC condition. The latter enabled to disentangle the global host transcriptional response within each subset and led to the identification of 20 distinct host gene expression profiles (one per cell type in each subset) (**Supp. Table 2**). Of note, we also calculated differential gene expression between NS1_R38A_ and WT subsets (e.g. NS1_R38A_ infected cells versus WT infected cells) and between infected and bystander subsets from the same condition (e.g. WT infected cells versus WT bystander cells). These comparisons and their results are summarized in Supplementary Tables (**Supp. Table 3**).

Compared to mock hAECs, we identified a combined total of 468, 153, 560, and 254 unique differentially expressed genes (DEG) in WT infected, WT bystander, NS1_R38A_ infected, and NS1_R38A_ infected subsets, respectively **(Fig. 5A**). Both common (i.e. present in all cell types) and cell type-specific DEGs were detected in each subset. Common DEGs that were upregulated in the WT and/or NS1_R38A_ infected subsets consisted mainly of genes related to the host antiviral response; however, more of these genes were upregulated in the NS1_R38A_ infected subset. Additionally, DEGs that were upregulated in both subsets were often induced to a higher amplitude in the NS1_R38A_ infected subset. For example, IFIT1 was significantly upregulated in secretory cells in both WT and NS1_R38A_ infected subsets; however, the increase was approximately 10-fold higher in the NS1_R38A_ infected secretory cells (**Supp. Table 3**).

**Figure 05.**
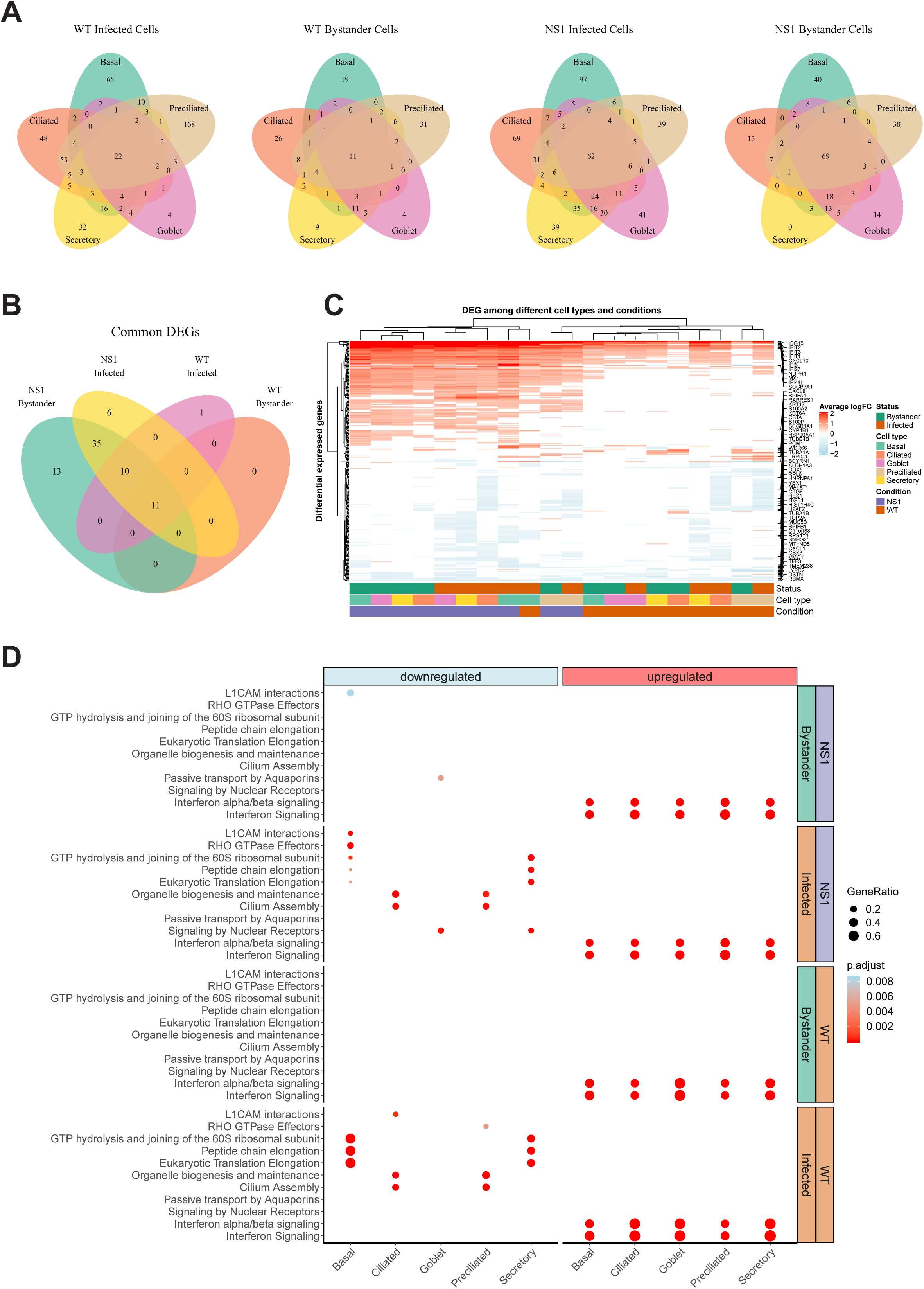
Global host antiviral response in WT and NS1_R38A_ virus-infected hAEC cultures. **A)** Venn diagrams showing the overlap of differentially expressed genes (DEG) among different cell types for each of the following: 1) WT infected cells, 2) WT bystander cells, 3) NS1_R38A_ infected cells, and 4) NS1_R38A_ bystander cells. For each comparison, DEGs identified in every cell type (“common” DEGs) are displayed in the centre of the Venn diagram. **B)** Venn diagram comparing the “common” DEGs identified in each of the comparisons above. A total of 10 core DEGs were present in all IAV-infected cells, regardless of the infection status (infected or bystander), cell type (ciliated, secretory, basal, goblet, or preciliated), or virus used for infection (WT or NS1_R38A_ virus). **C)** Hierarchical cluster analysis of DEGs identified in WT and NS1_R38A_ conditions among different cell types in both infected and bystander populations. For each of the 20 distinct DEG profiles identified, the top 5 upregulated (red) and downregulated (blue) DEGs are annotated in the heatmap. **D)** Dot plot illustrating pathway enrichment analysis performed on the 20 distinct DEG profiles. Enriched pathways are displayed on the left (y-axis) and the direction of enrichment is indicated at the top of the graph (downregulated in blue on the left panel; upregulated in red on the right panel). Significantly enriched pathways for WT (bottom 2 panels) and NS1_R38A_ (top 2 panels) are shown for infected (orange) and bystander (green) populations from each cell type (x-axis). Dots were adjusted in size and colour to illustrate the gene ratio and adjusted p-value for a particular pathway, respectively.

Clustering of the DEGs summarized in Figure 5A uncovered a core group of 10 genes that were consistently upregulated in WT and NS1_R38A_ cells, regardless of both infection status and cell type **(Fig. 5B**). This group included well-known ISGs (e.g. MX1, ISG15, and IFIT1), the transcription factor STAT1, genes involved in apoptosis (IFI27 and NUPR1), and IFI44L, which was recently identified as a feedback regulator for the host antiviral response ^66^. DEG clustering also revealed that on the whole, downregulated genes tended to be more cell type-specific **(Fig. 5C**). Interestingly, a number of downregulated DEGs were canonical cell type markers, including classical hAEC markers such as MUC5AC, MUC5B, ITGB1, and TUBB4B **(Fig. 5C**). Other classical markers were contra-regulated depending on the cell type. For example, SCGB1A1, a well-known marker for secretory cells, was significantly downregulated in NS1_R38A_ infected secretory cells, but significantly upregulated in WT infected basal cells **(Fig. 5C**). Finally, we identified multiple DEGs with established roles in cellular differentiation, proliferation, or migration **(Fig. 5C**).

To identify any significantly enriched biological pathways among the different conditions we next performed pathway enrichment analysis on the 20 distinct host gene expression profiles. This demonstrated that the majority of upregulated DEGs in both WT and NS1_R38A_ virus-infected hAECs were related to IFN signaling pathways **(Fig. 5D**). Notably, these pathways were not only enriched in all distinct cell types, but also in both the infected and bystander populations. More cell type-specific patterns were identified for downregulated DEGs, including depletion of the cilium assembly and the organelle biogenesis and maintenance pathways in both WT and NS1_R38A_ infected ciliated cells and preciliated cells (Fig. 5D). In addition, pathways associated with cap-dependent translation initiation were depleted only in WT and NS1_R38A_ infected basal and secretory cells. Finally, we also found that cell adhesion-associated pathways (L1CAM interactions) were depleted in both NS1_R38A_ infected and bystander basal cell populations, whereas Rho GTPase effector pathways were depleted in NS1_R38A_ infected basal cells only **(Fig. 5D)**. These data suggest that multiple cell types in the human airways, and particularly infected ciliated, basal, and secretory cells, undergo dynamic transcriptional changes following IAV infection that may alter the overall cellular composition of the respiratory epithelium. In addition, the depletion of these pathways may help explain the aforementioned differences in cellular composition we observed among mock, WT, and NS1R38A virus-infected hAECs in Fig. 3D.

### Important role for luminal cells in sensing and restricting incoming IAV

Given the complexity of the host antiviral response, as well as the paucity of information available on how distinct cell types in the human airway epithelium contribute to this response, we next examined the expression of multiple key antiviral signaling molecules in more detail. We aimed to establish a comprehensive map of the host antiviral response for each cell type during IAV infection. To achieve this aim, we first generated a manually curated list of genes related to essential innate immune and inflammatory pathways, including PRR genes, IFNs and their receptors, ISGs, as well as chemokines and cytokines (Table X). Cells were again grouped into 4 main subsets (WT infected, WT bystander, NS1_R38A_ infected, and NS1_R38A_ bystander) as in Figure 5.

As expected, many canonical host antiviral genes were strongly upregulated in the NS1_R38A_ virus-infected hAEC cultures, and to a lesser extent, in the WT virus-infected hAECs **(Fig. 6A**). This pattern was particularly evident for cytosolic PRRs (RIG-I/DDX58 and MDA5/IFIH1), the type III IFNs (IFNL1, IFNL2, and IFNL3), and most ISGs (e.g. IFIT1, IFIT2, IFIT3, ISG15). Interestingly, we found that in unexposed cells several genes were predominantly expressed in luminal cell types (ciliated, secretory, and goblet cells), including the endosomal PRR TLR3 and the transcription factor IRF1. Additionally, in unexposed cells, the PRR/adaptor gene STING/TMEM173 was mainly expressed in ciliated, preciliated, and basal cells, whereas the antiviral transcription factor IRF3 was ubiquitously expressed in all unexposed cell types **(Fig. 6A**). For genes with low basal expression levels, such as cytosolic PRRs RIG-I/DDX58 and MDA5/IFIH1 and transcription factors IRF7 and IRF9, no specific expression patterns were detected in unexposed hAEC cultures.

**Figure 06.**
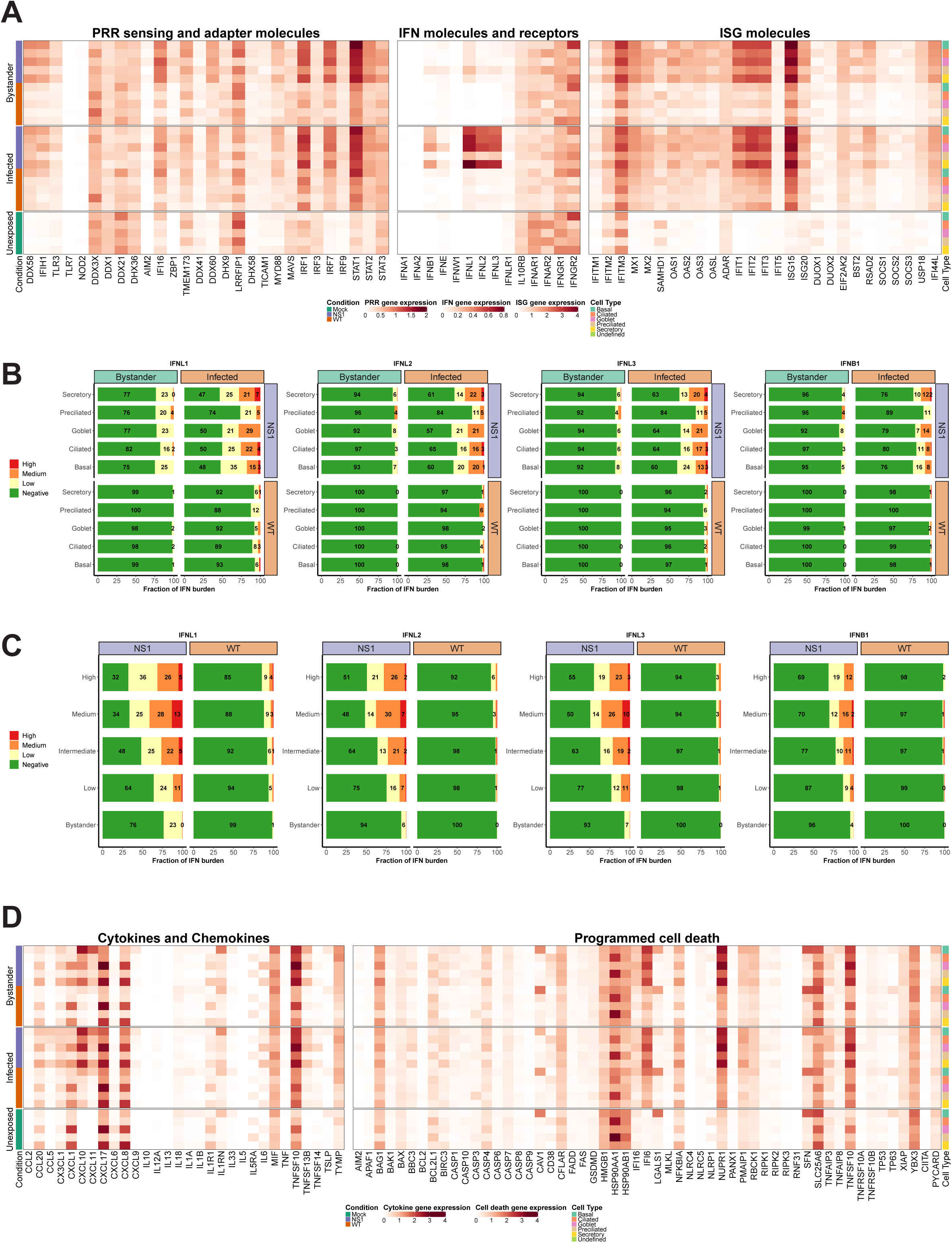
Cell type-specific host antiviral response to pandemic IAV infection. **A)** Heatmap illustrating the average expression levels for well-known antiviral genes, including PRR sensing and adapter genes (left panel), IFN genes and their receptors (middle panel), and ISG genes (right panel). Expression levels for individual genes are shown in columns and stratified by condition, infection status, and cell type (rows; representative colours shown in legends). **B)** Stacked bar plots showing the fraction of cells expressing low (yellow), medium (orange), or high (red) levels of either type I or III IFNs. Green represents the fraction of cells that were not expressing the IFN in question. For each bar graph, the cells are divided by condition, infection, status, and cell type. **C)** Bar graphs illustrating the fraction of cells expressing different levels of type I and III IFNs (low, medium, or high expression levels are coloured yellow, orange, or red, respectively; negative cells coloured green). In these plots, WT and NS1_R38A_ cells are divided by viral burden (i.e. bystander cell with no viral burden or infected cell with a low, intermediate, medium, or high viral burden; categories are labelled on the left side of each plot). **D)** Heatmap showing the average expression levels of various cytokines and chemokines (left panel) as well as many genes involved in programmed cell death (right panel). The heatmap is stratified by condition, infection status, and cell type (rows) for each gene (columns).

Following IAV infection, RIG-I/DDX58, MDA5/IFIH1, IRF1, and IRF7 expression levels were strongly induced, particularly in the NS1_R38A_ infected and bystander subsets. Expression of TLR3 and IRF9 was also upregulated, albeit to a lesser extent **(Fig. 6A**). For RIG-I/DDX58 and MDA5/IFIH1, we found that expression levels were highest in the NS1_R38A_ secretory, goblet, and basal cell populations in both infected and bystander subsets. Among the WT infected and bystander subsets, WT infected basal cells upregulated RIG-I/DDX58 and MDA5/IFIH1 expression to the greatest degree. IRF1 and IRF7 expression levels increased in most cell populations following IAV infection; however, IRF1 expression was still highest in luminal cell types. Notably, IRF3 expression was not upregulated following infection, likely because it is activated via a two-step phosphorylation event **(Fig. 6A**).

Type III IFNs were dramatically upregulated following IAV infection, especially in the NS1_R38A_ infected subset. IFNB1 expression was also elevated following infection; however, IFNL1, IFNL2, and IFNL3 expression clearly dominated the innate immune response. In contrast to IFNB1, other type I IFNs were not upregulated following IAV infection in hAECs **(Fig. 6A**). Interestingly, a recent scRNA-seq study in IAV-infected A549 cells also found that type III IFNs were highly induced after infection with a lab-adapted strain of IAV ^60^. When we systematically quantified the overall fraction of cells that expressed type I and/or type III IFNs in the WT and NS1_R38A_ virus-infected hAECs, we found that infected cells were the primary producers of both type I and type III IFN **(Fig. 6B**). Indeed for IFNL1, over 50% of cells in the NS1_R38A_ infected subset upregulated IFNL1 compared to 24% of cells in the NS1_R38A_ bystander subset. This pattern was also detected in the WT infected and bystander subsets, whereby 8% and 1% of cells upregulated IFNL1, respectively **(Fig. 6B**).

Within the WT infected subset, IFNL1 expression was fairly homogenous among distinct cell types; however, the exaggerated expression of IFNL1 in the NS1_R38A_ infected subset revealed that a greater fraction of ciliated, secretory, goblet, and basal cells upregulated IFNL1 compared to cells in the preciliated population **(Fig. 6B**). When cells were grouped according to the amplitude of IFNL1 they expressed (low, medium, or high), we found that the basal population contained a lower fraction of IFNL1-positive cells that expressed medium or high levels of IFNL1 compared to infected ciliated, secretory, or goblet cells. We also grouped cells by viral burden and analyzed the fraction of IFNL1-positive cells in the low, intermediate, medium, and high categories. Overall, cells with a lower viral burden also had a lower fraction of IFNL1-expressing cells. Despite this finding, we did not detect a significant correlation between viral burden and IFNL1 expression; however, the latter is likely due to the “high” IFNL1 category, which contained a smaller fraction of IFNL1-expressing cells than cells in the “medium” IFNL1 category (**Fig 6. B, Supp. Fig. 3**).

Interestingly, IFNL2 and IFNL3 displayed nearly identical expression patterns to IFNL1, but overall a lower fraction of cells expressed these IFNs and cells that did express them tended to do so at a lower amplitude (Fig. 6B, 6C). Of all IFNs, IFNB1 was upregulated in the smallest fraction of cells in all WT and NS1_R38A_ subsets. In the NS1_R38A_ infected subset, IFNB1 was expressed in 22% of cells across all cell types, whereas in the WT infected subset, IFNB1 was expressed in 2% of cells and was upregulated only in secretory, goblet, and basal cell types **(Fig. 6B**).

Multiple ISGs were strongly upregulated following pandemic IAV infection, including several that have been shown to inhibit various stages of the IAV life cycle. Similar to what we observed for PRRs and IFNs, ISG induction was most prominent in the NS1_R38A_ virus-infected hAEC cultures, but still strongly induced in the WT virus-infected hAECs **(Fig. 6A**). Notably, several ISGs were basally expressed in specific cell types in the unexposed (mock) hAEC cultures. For example, IFITM3, which was previously shown to restrict IAV entry into host cells, was basally expressed in secretory, goblet, and basal cell populations. Following IAV infection, IFITM3 expression was upregulated in most cell types in both WT and NS1_R38A_ virus-infected hAEC cultures. Similarly, we observed strong induction following IAV infection for the IFN-induced GTP-binding protein Mx1, the ubiquitin-like protein ISG15, and the cellular exonuclease ISG20 **(Fig. 6A**). Most IFIT family members were also highly induced in both WT and NS1_R38A_ virus-infected hAEC cultures, with the latter being most evident in secretory, goblet, and basal cells **(Fig. 6A**). Interestingly, the pattern and amplitude of SOCS1 expression, a negative feedback regulator, appeared to coincide with that of IFNL1. In contrast, we found that the negative feedback regulator USP18 was ubiquitously upregulated in all subsets **(Fig. 6A**). This indicates that virus-infected cells possibly modulate the autocrine Type I and III IFN signalling cascade.

On the whole, these results provide a comprehensive overview of the host IFN response to pandemic IAV infection in its natural target cells. They highlight an important role for luminal cells in sensing and restricting incoming respiratory viruses and identify several key antiviral genes that are induced in a cell type-specific manner following IAV infection. Finally, they demonstrate that infected cells are the primary producers of both type I and type III IFNs and that type III IFNs are the dominant IFNs driving the host antiviral response to IAV in human airway epithelial cells.

### High levels of inflammatory cytokines and chemokines in secretory and basal cells

Beyond the interferon response, IAV infection also activates the inflammatory response. The latter involves induction of multiple cytokines and chemokines, which in turn initiates immune cell recruitment from the bloodstream. This process, while crucial for viral clearance, can also exacerbate local inflammation and cause tissue damage. To better understand the host response to pandemic IAV in its natural target cells, we thus analysed the expression profile of key cytokines and chemokines in the unexposed, WT virus-infected, and NS1_R38A_ virus-infected hAECs. Of interest, we found that a number of cytokines and chemokines were basally expressed in unexposed secretory and goblet cells, including CCL20, CXCL1, CXCL17, and CXCL8. Conversely, other chemokines, such as CCL2, CXCL10, CCL5/RANTES, CXCL9, and CXCL11 were only detected in IAV-infected hAECs **(Fig.** XX).

Overall, we found that many inflammatory cytokines and chemokines were strongly upregulated following IAV infection, particularly in the NS1_R38A_ infected and bystander subsets. CXCL10, for example, was upregulated in most cell types in both WT and NS1_R38A_ virus-infected hAECs; however, its induction was especially prominent in NS1_R38A_ infected secretory, goblet, and basal cells. CXCL10 was also highly induced in NS1_R38A_ bystander secretory and basal cells, and to a lesser extent, in WT infected secretory and basal cells **(Fig. 6D**). A similar expression profile, albeit not as strong, was observed for CXCL11; however, in this case, we found that CXCL11 expression was highest in the basal cell population for all subsets. Interestingly, many cytokines and chemokines were expressed most prominently in basal cells. For example, CCL20 and CXCL9 expression was highly upregulated in NS1_R38A_ infected and bystander basal cells, whereas CXCL17 expression was strongly induced in WT infected basal cells and in NS1_R38A_ infected and bystander basal cells. CCL2 was also slightly upregulated in basal cells in NS1_R38A_ infected and bystander subsets **(Fig. 6D**). Finally, along with CCL2, CCL20, and CXCL9, CCL5/RANTES was barely detectable in WT infected and bystander subsets; however, it was upregulated in NS1_R38A_ infected secretory, ciliated, and basal cells.

IL6, a cytokine that has both pro-inflammatory and anti-inflammatory effects and has been linked to airway epithelial regeneration, was upregulated in secretory and goblet cells in both NS1_R38A_ infected and bystander subsets **(Fig. 6D**). Notably, IL13, a cytokine that can stimulate goblet cell differentiation and induce MUC5AC overexpression, was not upregulated in either WT or NS1_R38A_ virus-infected hAECs at 18 hpi ^67^. In contrast, macrophage migration inhibitory factor (MIF) expression was induced in most cell types in both WT and NS1_R38A_ bystander subsets. Lastly, we found that interleukin 1 receptor antagonist (IL1RN), a key modulator of IL1A and IL1B-related responses, was strongly upregulated in basal cells in both NS1_R38A_ infected and bystander subsets **(Fig. 6D**). Together these results suggest that at 18 hpi NS1_R38A_ virus-infected hAECs generate a much stronger inflammatory response to IAV infection than WT virus-infected hAECs. Moreover, upregulation of IL1RN indicates that NS1_R38A_ virus-infected hAECs may be trying to counteract this potent inflammatory response. Finally, our results show that the inflammatory response in both WT and NS1_R38A_ virus-infected hAECs is cell-type specific and that secretory and basal cells tend to induce the highest levels of inflammatory cytokines and chemokines, suggesting a critical role in bridging the innate and adaptive immune response.

### IAV infection induces a complementary programmed cell death (PCD) pathway response

Another important aspect of the inflammatory response is activation of programmed cell death (PCD) pathways. Recent studies have shown that in addition to inducing antiviral and inflammatory pathways, some PRRs can also activate PCD (ref). Moreover, IAV infection has specifically been shown to induce PCD pathways, such as apoptosis, necroptosis, and pyroptosis. For these reasons, and because many PRRs were strongly upregulated in WT and NS1_R38A_ virus-infected hAECs, we also determined the expression profiles of key genes involved in PCD in unexposed, WT, and NS1_R38A_ virus-infected hAECs.

Following IAV infection, we found that the pro-apoptotic death receptor ligand TNFSF10/TRAIL, which activates the cell extrinsic apoptosis pathway, was strongly upregulated in both WT and NS1_R38A_ virus-infected hAECs **(Fig. 6D**). Notably, in unexposed hAECs, TNFSF10/TRAIL was basally expressed in goblet and secretory cells; however, following infection its expression was upregulated in most cell types in WT and NS1_R38A_ virus-infected hAECs. This upregulation was particularly strong in the NS1_R38A_ infected and bystander subsets **(Fig. 6D**). Moreover, other members of the TNF family, including TNFSF13B/BAFF, TNFSF14/LIGHT, TNFAIP3/A20, and TNFAIP8, were also induced in the WT infected, NS1_R38A_ infected, and NS1_R38A_ bystander subsets. Interestingly, TNFSF13B/BAFF upregulation was strongest in secretory, goblet, and basal cells in both WT and NS1_R38A_ virus-infected hAECs, whereas TNFAIP3/A20 induction was highest in secretory, goblet, and ciliated cells **(Fig. 6D**).

Despite the strong upregulation of TNFSF10/TRAIL, other crucial effectors of extrinsic apoptosis remained unchanged following IAV infection. For example, we observed only a small increase in CASP3, CASP7, FAS, and TNFRSF10B/DR5 expression and no increase in CASP6, CASP8, TNFRSF10A/DR4, and FADD expression **(Fig. 6D**). Furthermore, several negative regulators of extrinsic apoptosis, such as CFLAR/FLIP and BIRC3/cIAP2, were upregulated following IAV infection. This upregulation was most prominent in the NS1_R38A_ infected and bystander subsets **(Fig. 6D**). A similar expression pattern was observed for key PCD genes involved in cell intrinsic apoptosis and necroptosis. For example, upon IAV infection expression of BAK, BCL2, MLKL, and RIPK3 remained unaltered. In addition, we found only a slight increase in BAX, RIPK1, and RIPK2 expression **(Fig. 6D**). However, the pro-apoptotic factors BBC3/PUMA and PMAIP1/NOXA were elevated in NS1_R38A_ infected and bystander subsets. Of interest, BBC3/PUMA expression was upregulated mainly in secretory and goblet cells, whereas PMAIP1/NOXA expression was prominent in NS1_R38A_ infected basal, secretory, and ciliated cells and NS1_R38A_ bystander basal cells **(Fig. 6D**). Intriguingly, we also found that the anti-apoptotic gene IFI6 was strongly induced in both WT and NS1_R38A_ virus-infected hAECs. Expression was upregulated in all cell types; however, it was highest in secretory, goblet, and basal cells in both WT and NS1_R38A_ virus-infected hAECs.

Finally, we found some expression changes in PCD genes involved in inflammasome activation and pyroptosis. For example, IFI16 and CASP1 expression levels were induced in NS1_R38A_ virus-infected hAECs, particularly in the NS1_R38A_ bystander subset. IFI16 induction was most prominent in goblet cells, whereas CASP1 expression was elevated in basal, secretory, and goblet cells. However, no expression changes were detected for PYCARD/ASC or GSDMD in either the WT or NS1_R38A_ virus-infected hAECs **(Fig. 6D**). Notably, NLRP3, which facilitates inflammasome activation in human macrophages, is not expressed in respiratory epithelial cells. These results demonstrate that at 18 hpi IAV infection induces a balanced duality between activation and repression of different PCD pathways.

### IAV infection leads to disruption of the airway epithelial cell barrier architecture

Because we observed diverse transcriptional changes related to cellular differentiation, proliferation, migration, and inflammation in the airway epithelium following IAV infection, we also assessed the gene expression signatures of host factors known to be involved in maintaining airway epithelial barrier integrity. The latter is essential to prevent severe epithelial damage and promote disease resolution. We first evaluated factors involved in cell adhesion, including several integrins and tight junction genes. Expression of tight junction protein 1 (TJP1/ZO1) was slightly decreased in both WT and NS1_R38A_ infected secretory cells. Additionally, the integrins ITGB1 and ITGAV exhibited reduced expression in NS1_R38A_ infected basal cells **(Fig. 7A**). Conversely, the tight junction protein CLDN4 was upregulated in NS1_R38A_ infected basal cells and in most cell types in the WT and NS1_R38A_ bystander subsets. We also found that expression of the pro-fibrotic factor IGFBP5 was increased in NS1_R38A_ infected and bystander secretory cells as well as in NS1_R38A_ infected goblet and basal cells **(Fig. 7A**).

**Figure 07.**
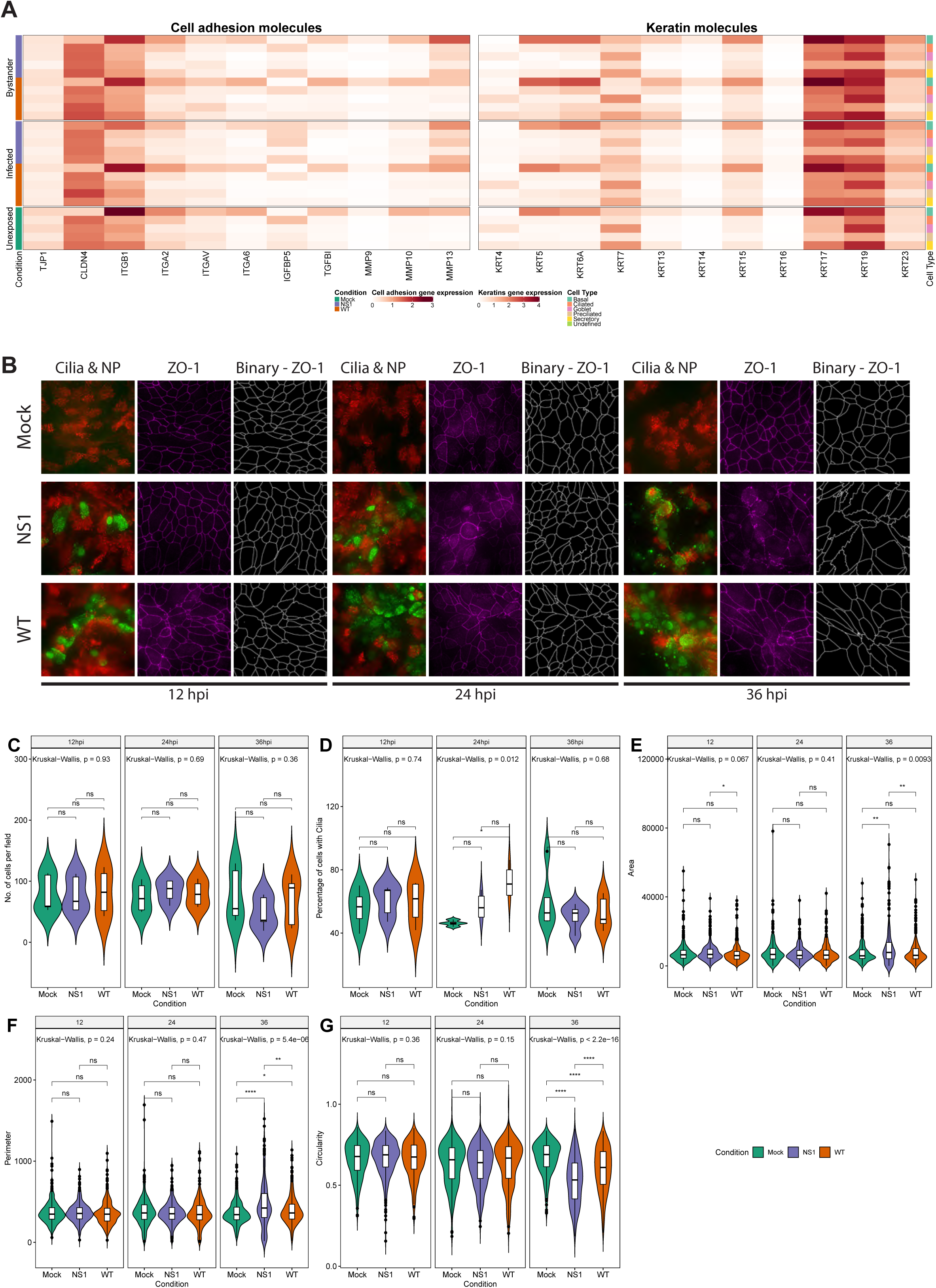
Disruption of the respiratory epithelium architecture by pandemic IAV infection. **A)** Heatmap illustrating the average expression levels of a variety of genes involved in cell adhesion (left panel) or keratin genes previously linked to cell differentiation in the respiratory epithelium (right panel). Expression levels for individual genes are shown in columns and divided by the condition, cell type, and infection status (rows). **B)** To monitor morphological changes hAECs were inoculated with 10,000 TCID_50_ of either WT or NS1_R38A_ IAV and fixed at 12, 24, and 36 hpi for immunofluorescence analysis. Formalin-fixed cultures were stained with different antibodies to highlight viral infected cells (nucleoprotein, green), ciliated cells (ß-Tubulin IV, red), and tight junction borders (ZO-1, purple). The tight junction images were binarized (ZO-1, white) using a custom image analysis script. Maximum intensity projection images obtained from z-stacks are shown for two different donors. Binarized tight junction images were used to calculate the following: **C)** the number of cells analysed overall, **D)** the percentage of cells with cilia, **E)** the surface area of individual cells, **F)** the length of the tight junction border of each cell (perimeter), and **G)** the shape of individual cells (circularity). Each measurement was calculated for mock (green), WT (orange), and NS1_R38A_ (purple) virus-infected hAEC cultures at 12, 24, and 36 hpi.

We also evaluated factors involved in cell migration, differentiation, and wound healing, such as members of the matrix metalloproteinase (MMP) and keratin families. Among the MMPs, MMP13 was upregulated in multiple cell types in the NS1_R38A_ infected and bystander subsets. In addition, we observed that MMP9 expression was slightly elevated in WT bystander basal cells, whereas MMP10 expression was reduced in NS1_R38A_ infected basal cells **(Fig. 7A**). The canonical basal cell marker KRT5 was upregulated in both WT and NS1_R38A_ bystander subsets and induced most prominently in the WT bystander basal cell population. Similarly, expression of KRT13 was upregulated in the WT bystander subset **(Fig. 7A**). Interestingly, KRT6A and KRT17, markers of progressive inflammation and wound healing, were also predominantly upregulated in the WT and NS1_R38A_ bystander subsets **(Fig. 7A**). Together these results indicate that pandemic IAV infection leads to dynamic, often cell-type specific, transcriptional changes in the airway epithelium that are known to have a detrimental influence on cell adhesion, barrier integrity, cell migration, differentiation, and wound healing.

Given the aforementioned transcriptional changes, as well as the virus-induced changes in cellular composition, we next examined how these changes influence the overall morphology of the airway epithelium. To this end, we performed a 36-hour time course experiment with mock, WT, and NS1_R38A_ virus-infected hAEC cultures. In this experiment, the hAEC cultures were fixed every 12 hours and then immunostained using antibodies against the viral NP protein, the tight junction protein TJP1/ZO1, and beta-tubulin 4 (TUBB4). TJP1/ZO1 and TUBB4 were used to assess the architecture of the epithelial barrier using z-stack images acquired over a distance of 25 - 30 microns **(Fig. 7B**). One feature we noticed was that in WT virus-infected hAECs the viral NP staining increased gradually over time, but in the NS1_R38A_ virus-infected hAECs the staining decreased from 24 hours onward **(Fig. 7B**). At 12 hpi, no gross morphological changes were observed in any of the hAEC cultures, which is in line with our previously results. However, after 24 hours we observed morphological aberrations in the TJP1/ZO1 tight junctions’ architecture in both WT and NS1_R38A_ virus-infected hAEC cultures. These aberrations were even more apparent at 36 hpi **(Fig. 7B**). Using the TJP1/ZO1 tight junction architecture as a reference, we quantified the morphology of individual cells at the different time points in more detail. Specifically, we used image-based analysis to measure cell surface area, circularity, and elongation of the cells **(Fig. 7C-G**). This revealed a slight reduction in the number of cells in the WT and NS1_R38A_ virus-infected hAEC cultures **(Fig. 7C**) that coincided with an increase in cell surface area, a loss of cell circularity, and an increase in the overall elongation of the cell periphery **(Fig. 7G**). Interestingly, we observed that these morphological aberrations occurred mostly in the proximity of the viral antigen-positive foci and that these changes were most pronounced in the NS1_R38A_ virus-infected hAEC cultures **(Fig. 7A**). Overall, these results indicate that the profound transcriptional changes in IAV-infected natural target cells at 18 hpi likely drive morphological changes in the human airway epithelium with a detrimental effect on airway epithelial morphology and likely on airway epithelial barrier integrity at 24 hpi and onwards.

## Discussion

To our knowledge, this is the first study to provide a comprehensive analysis of the cell type-specific host antiviral response to a respiratory virus infection in its natural target cells – namely, the human respiratory epithelium. Using scRNA-seq and primary well-differentiated hAEC cultures infected with either WT or NS1_R38A_ mutant IAV, we addressed multiple fundamental questions related to pandemic IAV infection in the human airways. Prior to our host transcriptional analysis, we first showed that all major cell types present in the human respiratory epithelium *in vivo*, could also be identified in primary well-differentiated hAEC cultures under both unexposed and virus-infected conditions. Moreover, in virus-infected hAECs, we demonstrated that both infected and bystander cells could be identified in each cell type. Interestingly, at 18 hpi, major changes in cellular composition were observed for both WT and NS1_R38A_ virus-infected hAECs compared to mock hAECs, including an overall decline in the number of ciliated and goblet cells as well as an increase in basal cell populations. Furthermore, we detected a shift in viral cell tropism from non-ciliated to ciliated cell types at later time points following IAV infection. Similar to previous studies, we found that the viral burden varied greatly among single infected cells, and additionally, we showed that this wide spectrum was present in all major epithelial cell types. An extensive analysis of the host antiviral response in both WT and NS1_R38A_ virus-infected hAECs, revealed 20 unique host transcriptional profiles (one for each cell type for both infected and bystander populations). As expected, NS1_R38A_-infected hAEC cultures induced a much stronger host innate immune response than WT-infected hAECs. Compared to bystander cells, we also found that infected cells were the main producers of IFNs, with a dominant role for IFN lambda. Additionally, we identified a number of cell type-specific differences in the host antiviral response, including a central role for luminal cells in sensing and restricting incoming IAV, and an important role for secretory and basal cells in terms of cytokine and chemokine production. Finally, we observed multiple changes in genes related to differentiation, proliferation, migration, and inflammation among these 20 different transcriptional profiles. Microscopic analysis of WT and NS1_R38A_ virus-infected hAECs at various time points following IAV infection suggested that the transcriptional changes we observed at 18 hpi were likely responsible for the downstream phenotypic changes in airway epithelial cell architecture during later stages of infection. Altogether, these results provide the first in-depth overview of the cell-type specific host antiviral response to pandemic IAV infection in the human airway epithelium.

Major cell types found in the human respiratory epithelium include basal cells, secretory cells, goblet cells, ciliated cells, and preciliated cells. In addition, rare cell types such as the newly identified ionocyte population, may also be present ^42,59,61^. Here, we used scRNA-seq to demonstrate that all major cell types found in the human respiratory epithelium *in vivo* can be annotated in primary well-differentiated hAEC cultures under both unexposed and virus-infected conditions. Moreover, in virus-infected conditions, we could identify infected and bystander cells for each major cell type. While we did identify several ionocytes in our analysis, due to the small number of cells a distinct cluster could not be identified. As such, we chose to exclude ionocytes from our subsequent analyses. However, it would be interesting to investigate whether ionocytes and other rare cell types contribute to the host antiviral response in future studies. Additionally, given the dynamic changes we observed in cellular composition at 18 hpi, it would also be interesting to analyze IAV-induced host responses at different stages of virus infection. However, due to technical interference by an increased abundance of viral mRNAs as well as the displacement of host and viral mRNAs into neighbouring single cell partitions, temporal analysis during viral infection may be challenging ^35,58^. Despite these challenges, our study provides a basic framework for future studies characterizing the host response to IAV and other respiratory pathogens in an authentic *in vitro* model that recapitulates the human respiratory epithelium *in vivo*.

Previous studies have demonstrated that at early time points of infection human IAV strains predominantly infect non-ciliated cells ^62^. Moreover, this tropism is thought to be due to the expression pattern of the 2,6-linked sialic acid receptor ^62,63^. We indeed found that at 6 hpi non-ciliated cell types were favoured by the human pandemic 2009 A/H1N1 IAV strain; however, at later time points the viral tropism changed to include other cell types, such as ciliated cells. The affinity of pandemic IAV for ciliated cells has been observed previously as early as 8 hpi – a finding that is consistent with our observations at 12 hpi and onwards ^68^. Here, we show for the first time, that secretory cells are the predominant target cells of human pandemic IAV and that goblet, basal, ciliated, and preciliated cells represent secondary target cells. This broad cell tropism may be driven by the relatively weak binding capacity of the viral HA protein to the 2,3-linked sialic acid receptor or by an alteration in sialic acid receptor distribution during infection due to changes in the overall cellular composition of the airway epithelium ^68,69^. Nonetheless, the broad cell tropism of human pandemic IAV towards cell populations generally targeted by avian-like IAV strains has the potential to facilitate the emergence of genetically reassorted novel IAV strains, some of which may have pandemic potential ^62^. Additionally, the finding that both non-ciliated and ciliated cells are infected by human pandemic IAV at different time points during infection may also be important in terms of IAV pathogenesis, disease progression, and future cell type-specific antiviral therapies.

In concordance with previous scRNA-seq studies using a lab-adapted strain of IAV, we also observed a wide spectrum of viral burden among cells infected with pandemic IAV ^35,36,60^. Our study also showed that the large heterogeneity in viral burden was not dependent on the intrinsic dsRNA binding capacity of the NS1 protein, as infected cells in WT and NS1_R38A_ hAEC cultures displayed a similar spectrum of viral burden. Moreover, since we detected this wide heterogeneity in viral burden among infected cells in all cell types, our results suggest that this phenomenon is not ascribed to a particular cell population. Other variables known to influence viral heterogeneity, such as defective interfering particles in the virus stock or the usage of a high MOI inoculum, were not present in our experimental settings ^33,35,70^. The former was corroborated by complete genome sequencing and the presence of viral mRNAs from all 8 segments of IAV in both WT and NS1_R38A_ hAEC cultures. Other variables like cell cycle state or expression levels of ISGs that can modulate virus replication did not show a strong positive correlation with viral burden. Thus, similar to previous studies in cell lines, the key factors that drive viral heterogeneity in primary well-differentiated hAEC cultures remain elusive; however, it is possible that this heterogeneity occurs stochastically ^33,35,70^. Future studies that simultaneous detect the host transcriptome and proteome within single virus-infected cells in a temporal setting may shed more light on this matter.

Our study underscores the importance of type III IFNs during IAV infection in the respiratory epithelium. Similar to Ramos and colleagues, we found that infected cells are the main producers of both type I and III IFNs and that IFN signalling occurs in both an autocrine and paracrine manner ^60^. Moreover, we found that IFNL1 was strongly induced following IAV infection in both WT and NS1_R38A_ virus-infected hAEC cultures and that IFNL2, IFNL3, and IFNB1 were induced to a lesser extent. We also provide a comprehensive overview of the innate immune response among distinct cell types in both infected and bystander cells following IAV infection. These analyses identified an important role for cell types that are exposed to the apical surface of the airway epithelium (luminal cells) in sensing and restricting incoming IAV. Luminal cells, including ciliated, secretory, and goblet cell types, were shown to be fully equipped to induce a robust IFN response towards an IAV inoculum with a relatively low MOI, which is in contrast to the previous observation in A549 cells ^35,60^. Although the reasons for these luminal cell differences remain unclear, they are likely driven by the fact that luminal cells are the first cells to encounter IAV and other respiratory viruses. In this context, it would thus be interesting to investigate whether a similar robust response ensues in these cell types after infection with another respiratory virus. Previous studies have shown that type III IFNs are produced prior to type I IFN upon IAV-infection, and that the amplitude of IFN production plays a pivotal role virus-induced pathogenesis _32,71_. We observed a much higher induction of IFN gene expression in NS1_R38A_ versus WT virus-infected cells, and in line with this we also observed a more pronounced change in the airway epithelium architecture in NS1_R38A_ virus-infected hAEC cultures. These results demonstrate that the hAEC culture model can be used to dissect the role of IFNs in virus-induced pathogenesis and respiratory epithelium barrier integrity.

Following IAV infection, we observed that several markers of progressive inflammation and wound healing were only upregulated in bystander cells, including several extracellular matrix modifying enzymes. Their transcriptional signature change at 18 hpi indicates disruption to the integral structural framework of the airway epithelial cell barrier and cellular homeostasis. This is in line with the observed deteriorated expression of cell type-specific markers in virus-infected cells and dynamic changes in the cellular composition at 18 hpi. Intriguingly, IAV-infected mice displayed similar deteriorated of cell type-specific marker expression in virus-infected cells ^32^. However, we demonstrate that the magnitude of the disruption to the airway epithelial cell barrier architecture coincides with the degree of the host immune response, as illustrated by the more pronounced disruption of the tight junction architecture in the NS1_R38A_ virus-infected hAEC culture. Resembling observed histopathologic changes in mice, rhesus macaques experiments, or humans that succumbed during the 1918, 1957, and 2009 influenza A virus pandemics ^29,72–75^. Thereby our study provides a novel framework to investigate the molecular mechanistic facets that underline ciliated epithelium degeneration and desquamation exterior of the adaptive immune response.

Combined these results for the first time highlight the Influenza A virus-induced dynamic and cell-type specific transcriptional changes that occur on a single cell level at the natural site of infection, namely the human respiratory epithelium. Therefore, this study embodies the first steps in generating a comprehensive overview of the complex virus – host interactions within the heterogenous cellular composition of the human respiratory epithelium.

## Acknowledgements

We would like to thank Prof. Dr. Martin Schwemmle, University of Freiburg, Germany, for providing the pdmH1N1 reverse genetic plasmids. This research was funded by the Swiss National Science Foundation grant numbers 179260 (RD), 160780 (VT), 173085 (VT), and the German Federal Ministry of Education and Research, project RAPID (VT).

