## Supplementary figures and images for "Comprehensive single cell analysis of pandemic influenza A virus infection in the human airways uncovers cell-type specific host transcriptional signatures relevant for disease progression and pathogenesis"

### Supplemental Figure 1

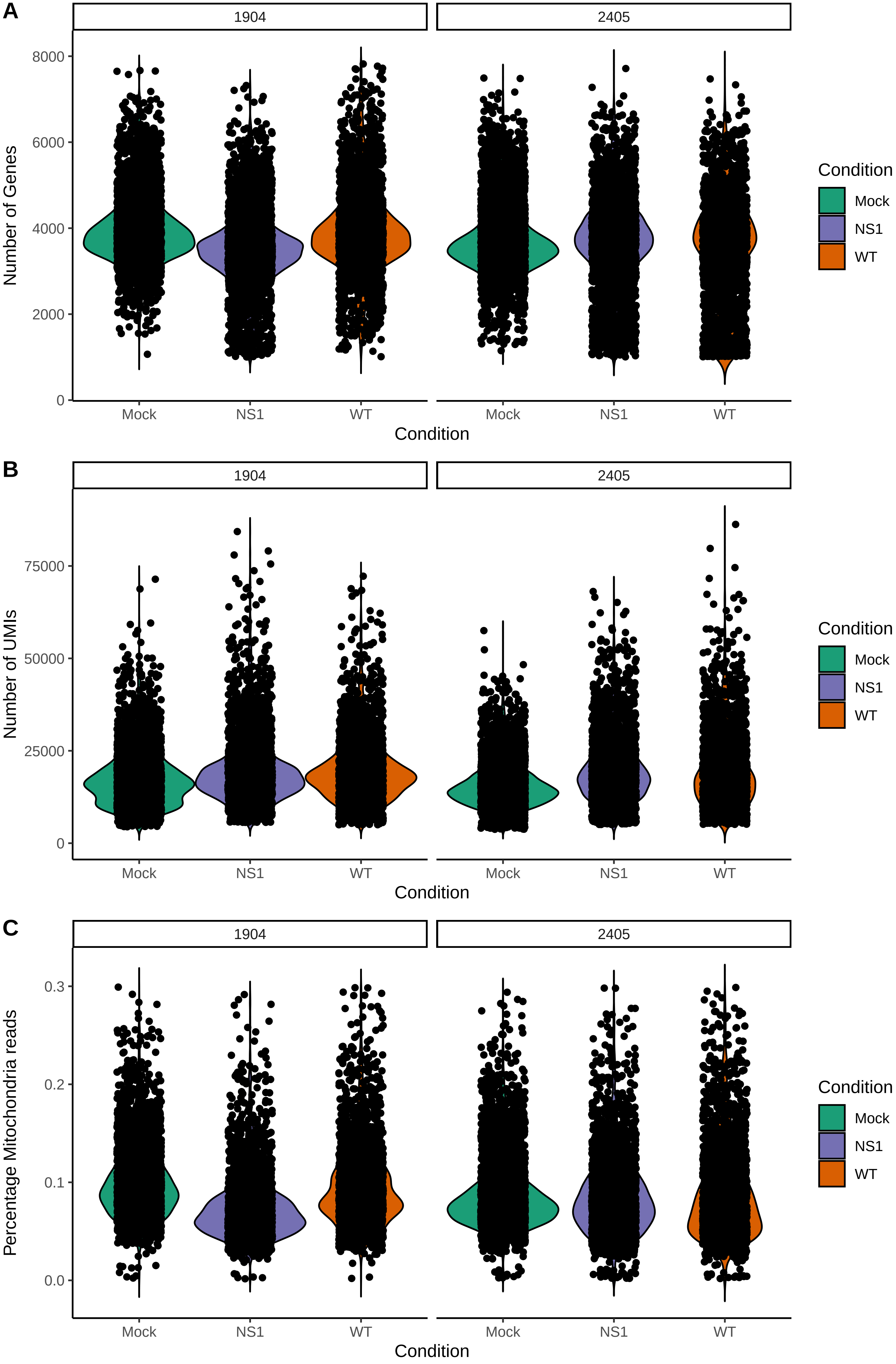

### Supplemental Figure 2

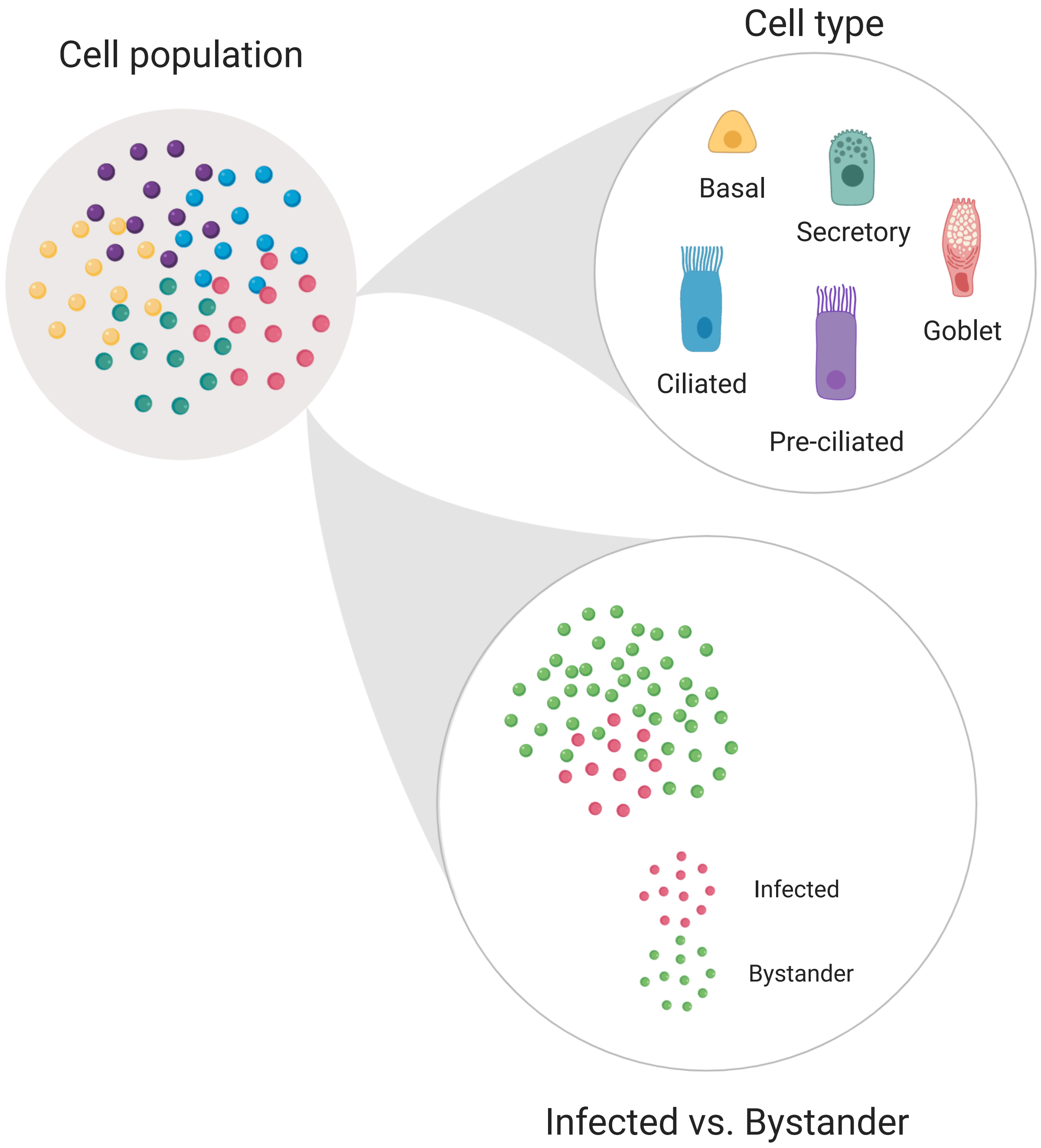

### Supplemental Figure 3

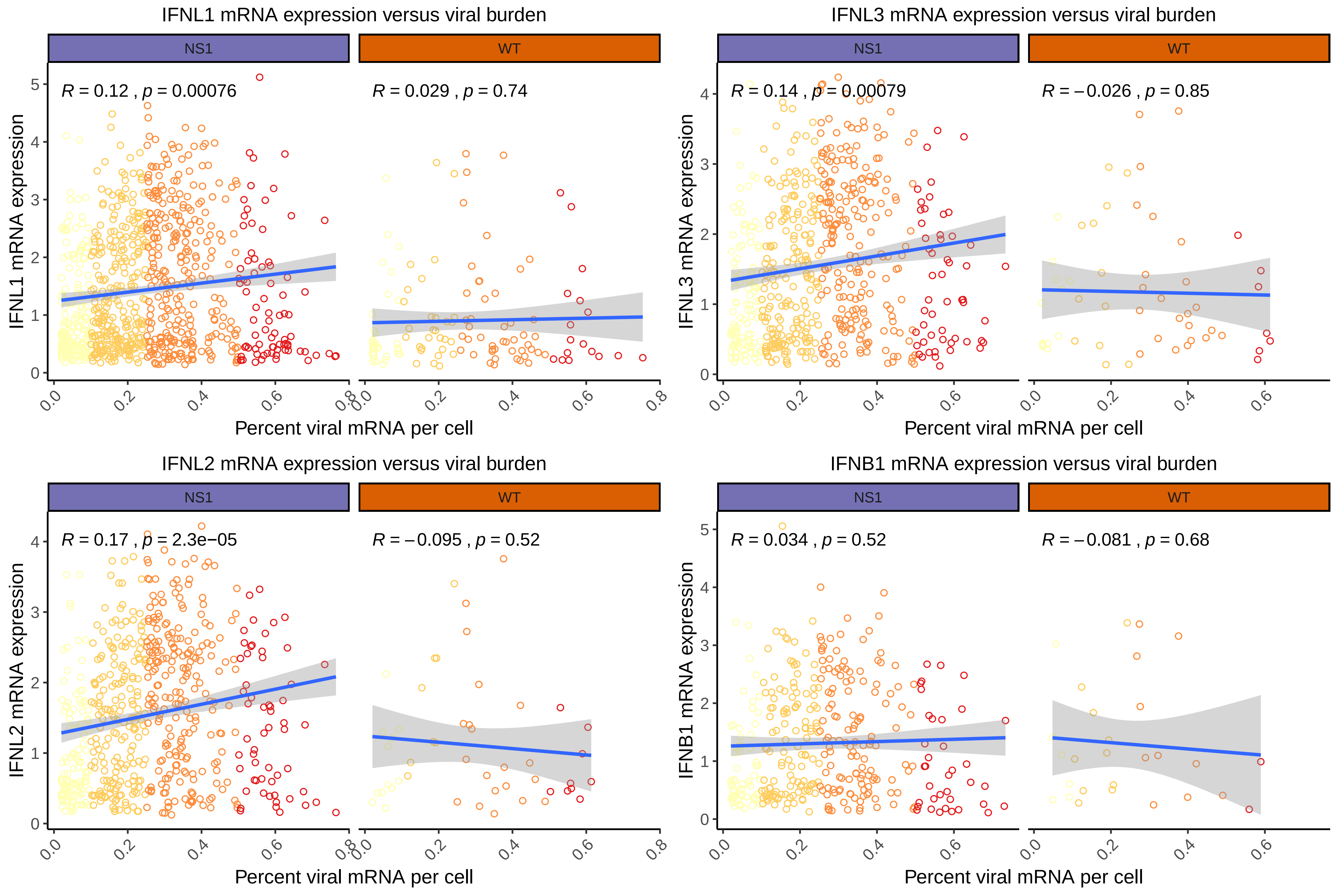
