## Supplementary Figure legends for "Comprehensive single cell analysis of pandemic influenza A virus infection in the human airways uncovers cell-type specific host transcriptional signatures relevant for disease progression and pathogenesis"

**Supplementary Figure 01. Visualization of quality control metrics for scRNA-seq data.** Data derived from each biological donor (1904 and 2405; top panels) for Mock (green), WT (orange), and NS1_R38A_ (purple) conditions were visually inspected using a variety of quality control metrics to select for high-quality cells prior to normalization and scaling. Violin graphs display **A)** the total number of genes, **B)** the total number of unique molecular identifiers (UMI), and **C)** the fraction of mitochondrial reads detected. Metrics are shown for single cells (black dots) in each condition (x-axis).

**Supplementary Figure 02.** **Schematic representation of different categorization variables.** For differential gene expression (DEG) analysis, single cells in each condition (Mock, WT, and NS1_R38A_) were further categorized based on their infection status (infected or bystander) and cell type (ciliated, secretory, basal, goblet, or preciliated). Cells were then placed in one of four main subsets for DEG analysis: 1) WT infected, 2) WT bystander, 3) NS1_R38A_ infected, or 4) NS1_R38A_ bystander subsets.

**Supplementary Figure 03. Correlation coefficient for interferon expression by viral burden.** To evaluate whether gene expression for interferon (IFN) lambda 1 (IFNL1), IFNL2, IFNL3, or IFN beta (IFNB1) correlated with viral burden, individual cells were ordered by the percentage of viral mRNA found in each cell (x-axis) and plotted against the corresponding gene expression values (y-axis) for both WT (orange; right panel) and NS1_R38A_ (purple; left panel) conditions. Individual cells are colored by viral burden as follows: low (≥2-10%; light yellow), intermediate (>10-25%; dark yellow), medium (>25-50%; orange) and high (>50%; red). Correlation coefficients for IFN gene expression and viral burden were calculated and are illustrated by a regression line (blue) with a 95%-confidence interval (grey). The Pearson correlation coefficient (*R*) and adjusted *p*-values are also shown for each gene.
